# Local G-quadruplexes are not a major determinant of altered gene expression in BLM-deficient cells

**DOI:** 10.1101/2023.09.08.556664

**Authors:** Thamar Jessurun Lobo, Peter M Lansdorp, Victor Guryev

## Abstract

Mutations in the *Blm* gene can cause Bloom Syndrome, a genetic disorder characterized by genome instability and cancer predisposition. *Blm* encodes a helicase which was reported to resolve G-quadruplex DNA structures *in vitro*. The G-quadruplex resolving activity of the BLM helicase has been previously implicated in altering gene expression. However, the exact mechanisms of how G-quadruplex structures may affect gene expression remain to be elucidated. We employed experimentally defined G-quadruplex forming DNA sequences and generated transcriptomes for several Bloom Syndrome patient-derived cell lines and BLM-deficient mouse embryonic stem cells to further investigate the effect of G-quadruplexes on gene expression. Our results do not support the previous findings that G-quadruplexes located within a gene play a major role in altering its expression in BLM-deficient cells. We found concerted large-scale changes in transcript abundance, splicing, nucleosome occupancy, and phasing, that cannot be linked to the local presence of G-quadruplex sequences in either gene bodies or promotors. The investigation of genomic features associated with large-scale differences in nucleosome density highlights the rDNA locus and active enhancers as the most strongly affected regions. We hypothesize that global changes in chromatin structure rather than local G4s might mediate the transcriptome changes in the absence of BLM.

## Introduction

Bloom Syndrome (BS) is a rare disorder characterized by growth deficiency, increased sensitivity to sunlight, moderate immune deficiency, and decreased fertility. Due to a high probability of developing multiple types of cancers, BS patients have a strongly reduced life expectancy. The underlying cause of these symptoms is a mutated gene, which encodes for the BLM helicase (1). BLM is a DNA helicase that unwinds double-stranded DNA in a 3’-5’ direction (2). It plays an important role in DNA replication and is recruited to stalled replication forks. BLM has mainly been implicated in the prevention and repair of double-strand breaks by homologous recombination, as reflected by the high rates of sister chromatid exchanges (SCEs) in BS patient cells (3, 4, 5, 6, 7). In situ hybridizations localize BLM to nucleolus, telomeres and nuclear PML bodies (22), suggesting a role in preservation of genome stability in regions that are difficult to replicate. The BLM helicase is also known for its high affinity for G-quadruplex (G4) structures, which it is reportedly able to resolve *in vitro* (8). G4s are stacked DNA or RNA structures that have long been suspected to have a biological function with suggested roles in telomere maintenance, replication, and transcription (9). Naturally, BLM’s G4-resolving abilities have raised the question of whether BS symptoms might be a consequence of dysregulated G4 formation. On the transcription level, a previous study reported that altered gene expression in BS fibroblasts was associated with the presence of sequence motifs having G-quadruplex forming potential, on either strand up– and downstream of the transcription start site (TSS), in the first three introns, and up– and downstream the transcription termination site (TTS) of those genes (10). A later study profiled the transcriptome of fourteen BS fibroblast cell lines and twelve controls, including the cell lines profiled by the aforementioned study. The authors reported that BS-specific transcriptional differences were correlated with the presence of G4 motifs on either strand in the first intron and, to a lesser extent, on the transcribed strand up– and downstream of the TSS (11).

As the formation of G4s has been difficult to examine experimentally, both studies made use of *in silico* predicted G4 motif locations, such as canonical G4 motifs with 1-7 bp loops (G_3+_N_1–7_G_3+_N_1−7_G_3+_N_1−7_G_3+_) (10) and its variant with extended loops of up to 12 nucleotides (11). However, the set of potential G-quadruplex structures was recently greatly expanded by a new sequencing-based method resulting in a genomic map of experimentally defined G4 forming DNA sequences in human genomes (12, 13). This map enables a more comprehensive investigation of the role of G4s in BS.

Because G4 sequences are typically GC-rich, it is important to test whether the transcription in BS cells is primarily affected by the G4 forming ability or the higher duplex stability of GC-rich genomic regions. The previous studies tested for the effect of GC-richness indirectly, either by comparing the GC content between differentially expressed genes and a random set (11) or checking the enrichment of imperfect G4 motifs with a disrupted or absent G-run (10).

We expanded the existing microarray-based expression data by profiling transcriptomes of human BS human lymphocyte and fibroblast cell lines, and F1 hybrid murine embryonic stem (ES) cells with *Blm* knock-out, using RNA-seq. In our study, we investigated the relationship between local experimentally defined G4 forming DNA sequences and BLM deficiency-specific transcriptional alterations for each dataset. We employed linear regression to enable explicit correction for local GC content and to omit the use of arbitrary cut-offs for differential expression. As an alternative approach that is unaffected by technical biases, we also performed an analysis of allele-specific expression in the F1 hybrid murine cells investigating the expression of genes discordant for the presence of G4 motifs between parental haplotypes. Next to the expression levels of genes, we also investigated the chromosomal co-localization of genes dysregulated upon BLM deficiency as well as its role in alternative splicing.

## Materials and Methods

### Reanalysis of microarray expression profiles

Original CEL files were downloaded from GEO, accession GSE54502. The dataset was processed and normalized using Transcriptome Analysis Console (ThermoFisher Scientific) version 4.0.2 using default settings. We compared BS and control groups taking age and gender as additional factors and used the eBayes method for differential expression analysis.

### Cell cultures

The cell lines from Bloom Syndrome and control patients were obtained from the Cell Repository of Coriell Institute: GM07492 and GM07545 (primary fibroblasts, control), GM02085 and GM03402 (primary fibroblasts, BS), GM12892 (EBV-transformed B-lymphocytes, control), GM16375 and GM17361 (EBV-transformed B-lymphocytes, BS). Mouse ES cell line F121.6 (F1 hybrid between 129/Sv and Cast/EiJ) was a kind gift from Prof. J. Gribnau (Erasmus MC, Rotterdam, The Netherlands). *Blm* mutant ES cell lines were generated as previously described (7). Fibroblasts were cultured in Dulbecco’s modified Eagle’s medium (DMEM) (Life Technologies) supplemented with 10% v/v fetal bovine serum (FBS) (Sigma Aldrich) and 1% v/v penicillin-streptomycin (Life Technologies), B-lymphocytes in RPMI1640 (Life Technologies) supplemented with 15% v/v FBS and 1% v/v penicillin–streptomycin. ES cells were cultured on mitotically arrested mouse embryonic fibroblast cells in DMEM (Life Technologies), supplemented with 15% v/v FBS (Bodinco BV), 1% v/v penicillin-streptomycin, 1% v/v non-essential amino acids (Life Technologies), 50 µM 2-mercaptoethanol (ThermoFisher Scientific), and 1,000 U ml^−1^ leukemia inhibitor factor (Merck). All cells were cultured at 37 °C in 5% CO_2_.

### RNA-seq library preparation

Exponentially growing cells were harvested from individual cultures for each sample, and RNA was isolated using the Nucleospin RNA kit (Macherey Nagel). Libraries were generated for each of the three replicates, with exception of one sample where isolation produced RNA with a low RIN score (below 7) and thus was done in duplicate. RNA-sequencing libraries were prepared using the NEBNext Ultra RNA Library Prep kit for Illumina (NEB) combined with the NEBNext rRNA Depletion kit (NEB). Complementary DNA quality and concentrations were assessed using the High Sensitivity dsDNA kit (Agilent) on the Agilent 2100 Bio-Analyzer and the Qubit 2.0 Fluorometer (Life Technologies).

### Illumina sequencing

Clusters were generated on the cBot (HiSeq2500) and single-end 50 bp reads were generated using the HiSeq2500 sequencing platform (Illumina).

### Preprocessing and analysis of RNA-seq expression data

Quality trimming of raw reads was performed using single-end mode implemented in Trimmomatic package (v. 0.36) using recommended settings and a minimum read length after trimming of 20 bp. Reads were mapped to the primary human genome assembly (GRCh38) or mouse genome (GRCm38) and quantified with STAR aligner (v. 2.7.0f) using respectively release 92 of the human gene annotation and release 84 of the mouse gene annotation from Ensembl (http://www.ensembl.org).

Genes with an average expression of <= 1 FPM were excluded from the analysis. Using the R package EdgeR (v. 3.26.8) a quasi-likelihood negative binomial generalized log-linear model was fitted to estimate differential expression between controls (reference) and BS/*Blm* mutation for each cell type (while correcting for gender in the case of the fibroblast dataset and ignoring non-autosomes downstream for the lymphocytes in downstream analysis, as we could not correct for gender). Transcriptional differences between replicates of GM07492 (reference) and GM07545 were estimated in the same manner. For this set, genes on non-autosomes were ignored in downstream analysis, as we could not correct for sex-specific differences in our analysis.

### Analysis of allele-specific expression

For quantification of allele-specific expression, we extracted variants where both genotypes were homozygous and supported by at least 5 sequencing reads, and discordant between 129 and CAST mice. We used variants reported by the mouse genome project (http://www.sanger.ac.uk/data/mouse-genomes-project version 6). To ensure the specificity of allele calling we employed the WASP function of STAR aligner and extracted strain-informative reads for 129 and CAST genetic backgrounds into separate alignment files. We used htseq v 0.6.1.p1 for the quantification of allele-specific expression.

The differential allele-specific expression between 129/Sv and Cast/EiJ backgrounds in *Blm* mutant cells and controls was also estimated using the EdgeR package by fitting a quasi-likelihood negative binomial generalized log-linear model, after exclusion of lowly expressed genes (average expression of <= 1 FPM). Genes on non-autosomes were ignored in downstream analysis.

### Definition of the primary transcript and gene partitions

For annotation of promotor, intron flanks, and other regions we first determined the primary transcript for each expressed gene. We performed guided transcriptome annotation using Stringtie (v.2.1.5, options –G –e) per each of the three RNA-seq datasets (human lymphocytes, human fibroblasts, mouse ES cells). For each gene, we selected the transcript with the highest fragments per million per kilobase (FPKM) value as the primary transcript.

The following regions were used: Upstream TSS (250 bp immediately upstream of the transcription start site of the primary transcript, TSS); Downstream TSS (250 bp immediately downstream TSS); Introns 1, 2, and 3 (250 bp immediately downstream 5’-end of the intron), Upstream TTS (250 bp immediately upstream the transcription terminations site of primary transcript; and Downstream TTS (250 bp immediately downstream the transcription terminations site of primary transcript). The distances (250bp) were selected to match the definitions of the previous paper (11).

### Regression analysis

Locations of experimentally defined G4 forming DNA sequences in the human and mouse genome in K^+^ conditions were downloaded from GEO database, accession GSE110582 in BED format. Base mismatch percentages for both the human and mouse genome were downloaded from the same accession as BEDgraph files. Coordinates were lifted from hg19 to hg38 assembly using the UCSC lift-over genome annotation tool with default settings. For each selected region the GC percentage (pGC) was retrieved and the G4 sequences count (nG4) was quantified for both the coding and template strand by counting the number of G4 forming DNA sequences overlapping the region at each strand. In addition, we calculated “G4scores” for both the coding and template strand at each selected gene region by unweighted averaging the mismatch percentage as reported in the BEDgraph files for all genomic regions overlapping the gene region using the map command from bedtools v2.17.0.

For the identification of G4 motifs discordant between 129 and CAST mice, we employed a sliding window search (window size = 100bp, using GRCm38 reference and 1bp step), applying strain-specific allele and checking for the presence of a G4 motif with 4 G tracks of 3 or more bases with loop sizes 1-7bp. Non-redundant G4 motifs only seen in one mouse strain were reported as discordant. GC content for each partition of a gene was calculated based on GRCm38 coordinates by applying 129/CAST alleles before calculating sequence GC percentage.

Transcriptional changes (ΔT) were characterized as –ln(P-value) multiplied by the sign of the fold change, meaning that positive and negative values indicate up and downregulation in the Blm samples, respectively. To identify the effects of respectively the presence of G4 sequences, G4 sequence count, and G4-score on transcription, the linear model Δ*T* ∼ *x_upstream TSS_* + *x_downstream TSS_* + *x_intron_*_1_ + *x_intron_*_2_ + *x_intron_*_3_ + *x_upstream TTS_* + *x_downstream TTS_* was fitted with *x =* (*nG*4 ≠ 0), *x* = *nG*4 and *x* = *G*4*score* for both coding and template strands. To identify an effect of GC percentage, the linear model Δ*T* ∼ *pGC_upstream TSS_* + *pGC_downstream TSS_* + *pGC_intron_*_1_ + *pGC_intron_*_2_ + *pGC_intron_*_3_ + *pGC_upstream TTS_* + *pGC_downstream TTS_* was fitted. To determine the relative importance of G4 motifs and the GC percentage, we fitted the model Δ*T* ∼ *x_upstream TSS_* + *x_downstream TSS_* + *x_intron_*_1_ + *x_intron_*_2_ + *x_intron_*_3_ + *x_upstream TTS_* + *x_downstream TTS_* + *pGC_upstream TSS_* + *pGC_downstream TSS_* + *pGC_intron_*_1_ + *pGC_intron_*_2_ + *pGC_intron_*_3 +_ *pGC_upstream TTS_* + *pGC_downstream TTS_* with *x =* (*nG*4 ≠ 0), *x* = *nG*4 and *x* = *G*4*score* for both coding and template strands. We also fitted the models above separately for only up– and only downregulated genes. In this case, ΔT was just defined as –ln(P-value) for each set.

Allele-specific transcriptional changes (ΔASE) were characterized as –ln(P-value) with positive and negative values indicating up and downregulation of the Cast allele expression relative to that of the 129 strain. G4 motif discordance (ΔG4) was characterized as respectively –1 and 1 when the Cast allele contained less or more G4 motifs than the 129 allele, or 0 when there was no difference. To identify an effect of G4 motif discordance, the linear model Δ*ASE* ∼ Δ*G*4 *_upstream TSS_* + Δ*G*4*_downstream TSS_* + Δ*G*4*_intron_*_1_ + Δ*G*4*_intron_*_2_ + Δ*G*4*_intron_*_3_ + Δ*G*4 *_upstream TTS_* + Δ*G*4*_downstream TTS_* was fitted for both coding and template strands. To identify an effect of the average GC percentage of the two alleles, the linear model Δ*ASE* ∼ 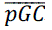*_upstream TSS_* + 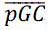*_downstream TSS_* + 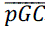*_intron_*_1_ + 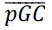*_intron_*_2_ + 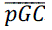*_intron_*_3_ + 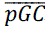*_upstream TTS_* + 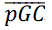*_downstream TTS_* was fitted. To determine the relative importance of G4 motif discordance and the average GC content, we fitted the linear Δ*ASE* ∼ Δ*G*4 *_upstream TSS_* + Δ*G*4*_downstream TSS_* + Δ*G*4*_intron_*_1_ + Δ*G*4*_intron_*_2_ + Δ*G*4*_intron_*_3_ + Δ*G*4 *_upstream TTS_* + Δ*G*4*_downstream TTS_* + 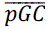*_upstream TSS_* + 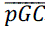*_downstream TSS_* + 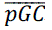*_intron_*_1_ + 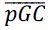*_intron_*_2_ + 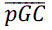*_intron_*_3_ + 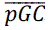*_upstream TTS_* + 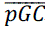*_downstream TTS_* for both coding and template strands.

### Analysis of the influence of Blm knock out on expression of alleles discordant for G4

We selected genes that have discordant G4 motifs between 129 and CAST genetic backgrounds and overlapped one of the gene partitions mentioned above. We required that one of the alleles complied with G4 motif (G_3+_ N_1-7_ G_3+_ N_1-7_ G_3+_ N_1-7_ G_3+_), while the same region in the other genetic background did not match this formula due to SNV or indel affecting one of the G-stretches. We treated G4 motifs on coding and template strands of the transcripts separately. We discarded regions that contained multiple discordant G4 motifs with opposite distribution (e.g., intron 1 with a G4 motif that is present in 129 and not CAST version of the locus, when the same intron also harbored another G4 motif on the same strand that is present in CAST, but not 129 genetic background). We called allele-specific expression for 129 and CAST alleles in combined wild type and combined *Blm* knock-out samples as mentioned above. We next discarded lowly expressed genes with allelic expression below 1 count per million (CPM). We calculated the expected expression of each G4-allele in the *Blm* knock out as:

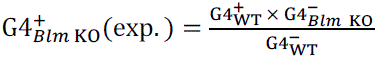

We then determined log_2_ fold change between observed and expected expression of G4-alleles in the *Blm* knockout background per partition per coding/template strands. We used a two-tailed T-test to check if the distribution of log_2_FC is significantly different from zero. Significantly positive values would indicate common upregulation of G4-allele in *Blm* –/– background, while negative – downregulation. We also performed a one-tailed F-test to test if the ratios show higher variability compared to genes expressed above 1 CPM level, but containing no discordant G4 motifs in any of the gene partitions considered in our analysis.

### Analysis of alternative splicing

Alternative splicing analysis was performed using rMATS-turbo v 4.1.0 (14) with default settings and the same alignment files (BAM format) used for differential expression analysis. Splicing events with FDR below 0.01 were considered significant. We used a two-tailed *X*^2^ test to check if significant splicing events were unevenly distributed among skipping and retention events. For example, if the share of significant exon skipping (from all exon skipping events) was higher than the share of significant exon retention events. The same *X*^2^ test was used for checking if introns containing a G4 motif (loop size 1-7bp) in their first 250 bp are more likely to be among significant splicing events. Introns following skipped, upstream and downstream exons were tested.

### Clustering of differentially expressed genes

Differentially expressed genes (FDR cutoff 0.01) were grouped into closest neighbor pairs based on their genomic coordinates. Gene pairs with intergenic distances above 100 kbp were removed. For the remaining pairs, the directions of change were compared and recorded. We used a two-tailed *X*^2^ test of independence to compare the total number of observed numbers of neighboring gene pairs with the same and opposite directions of expression change versus the expected number of pairs with the same and opposite directions, given the proportion of up– and down-regulated genes.

### Visualization of differentially expressed genes

For visualization we used genes that were significant (FDR cutoff 0.01) in at least one of the following sets: microarray fibroblast dataset, RNA-seq fibroblast dataset, and RNA-seq lymphocyte dataset. Normalized expression data – RMA values for the microarray dataset and FPM values for the RNA-seq dataset were converted to z-scores and plotted using pheatmap R package.

### Gene Ontology enrichment analysis

First, Ensembl gene identifiers were converted to Entrez gene identifiers. Only genes with one-to-one matches were kept. We used the *goana* function from the R package Limma (v. 3.46.0) with an FDR cutoff of 0.1 to perform Gene Ontology enrichment analysis for the microarray expression profiles. For the RNA-seq expression profiles, we used the *goana* function from the R package EdgeR with an FDR cutoff of 0.01.

### Analysis of relative nucleosome occupancy

We utilized data from Strand-seq libraries from wildtype and *Blm* –/– mouse ES cells which were obtained by MNase digestion of DNA (7), sequenced in paired-end mode (2×105bp), and are available through ArrayExpress accession E-MTAB-5976. We merged alignments for all wildtype libraries to use in the quantification of nucleosome occupancy of WT sample. We merged BAM files of all *Blm* –/– Strand-seq libraries to represent nucleosome profiles of *Blm* knockout cells. We processed the two alignments discarding unmapped, PCR duplicates, secondary alignments, QC failures and reads with template lengths below 100bp or above 220bp. We focused on unambiguously mapped reads (mapping quality of 20 or higher). For analysis of relative nucleosome occupancy for gene partition sets, G4 sequence, and G4 motif we used the same definition of these regions as described above. CpG island profile was downloaded from UCSC genome browser. For nucleosome phasing analysis, we used properly paired read pairs that fulfill the same filtering as described above. We determined the center of each nucleosome as a rounded median point between start and end of a fragment and quantified the number of fragment pairs that show distance between their center in the range 150-500bp.

## Results

### Correction for GC content eliminates the relationship between the presence of local G4 sequences and BS-specific transcriptional changes seen in microarray data

First, we employed linear regression to test whether the local presence of experimentally defined G4 forming DNA sequences, similar to predicted G4 motifs, shows significant overrepresentation in genes differentially regulated in BS cells. Since one of the previous studies (10) used a subset of the BS fibroblast cell lines that were transcriptionally profiled by the other study (11) we chose to reanalyze the microarray data generated by the latter study. We found that the presence of G4 sequences (Figure 1A) on either coding or template strand at the start of the second and third intron and up– and downstream of the TTS were correlated to differential expression (p-value < 0.01). Interestingly, GC content also showed a significant correlation to BS-specific alteration of gene expression in all gene partitions but downstream of the TSS and TTS (Figure 1A). Since both the local presence of G4 sequences and the GC content were significantly correlated with transcriptional changes, we tested which was more important by including both variables into the same regression model (Figure 1). We found that, by taking GC content into account, the predictive power of local G4 sequences diminished greatly: it was no longer significant in any of the gene partitions. However, the predictive power of GC content remained unaffected by the addition of G4 sequences. To illustrate this effect, Figure 1B displays the connection between BS-specific transcriptome alterations and the GC content and presence of G4 sequences on the template strand around 3’-UTR (upstream of the TTS).

**Figure 1.**
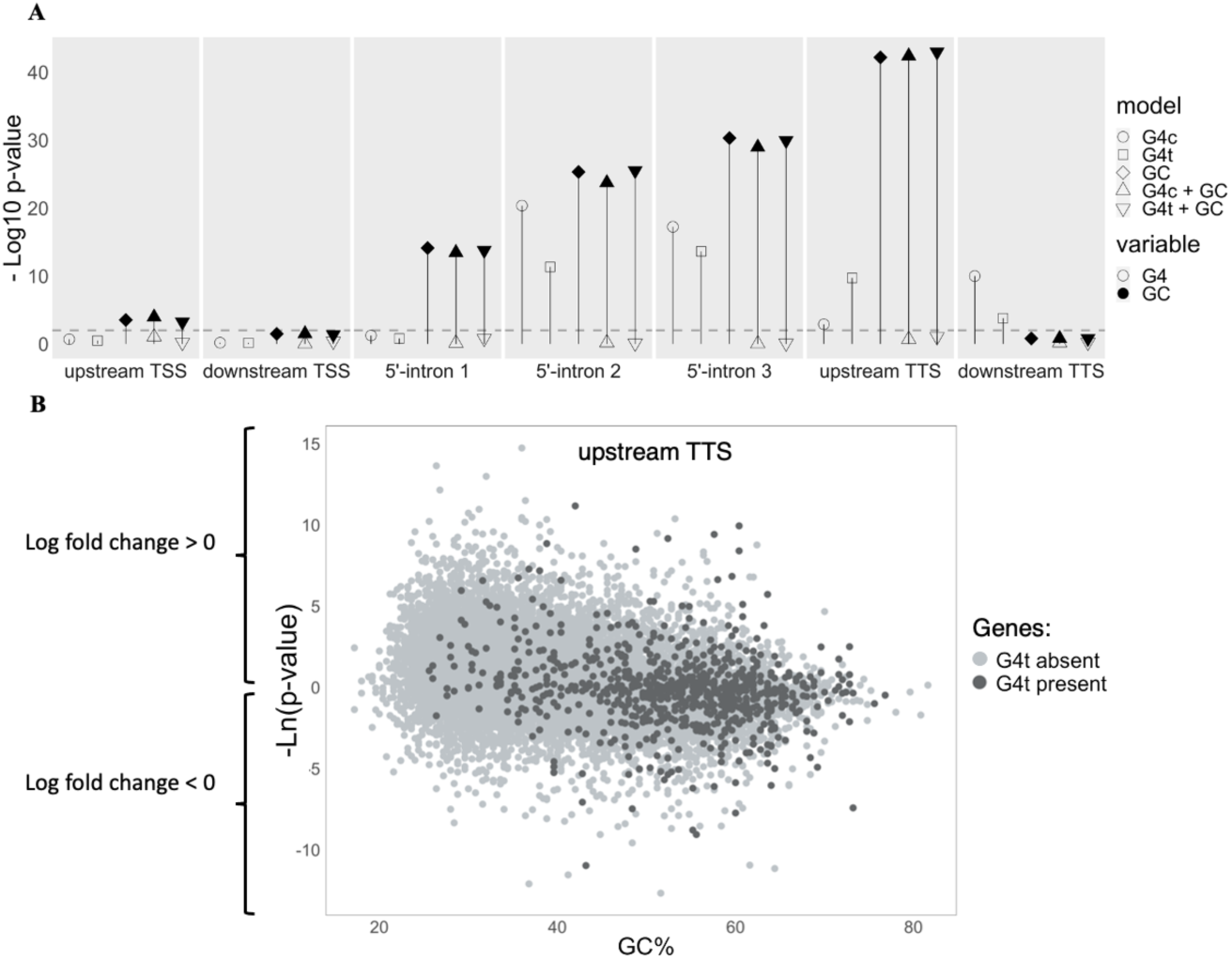
Results of regression analyses for BS-specific transcriptome alterations in microarray data of fibroblasts. (A). –log_10_ of the p-value for each variable, indicating the strength of the association of the presence of experimentally confirmed G4-forming sequences (no fill) on either the template (G4t) or coding strand (G4c) and GC content (black fill) in various gene regions with the transcriptome alterations between BS and control fibroblasts. The shape of the data points indicates the regression model: the G4 presence on the coding strand of each gene region (G4c, circle), the G4 presence on the template strand of each gene region (G4t, square), the GC content of each gene region (GC, diamond), both the G4 presence on the coding strand and the GC content of each gene region (G4c + GC, triangle pointing up) or both the G4 presence on the template strand and the GC content of each gene region (G4t + GC, triangle pointing down). The dashed grey line marks p-value=0.01. (B). Scatterplot showing BS-specific transcriptome alterations and the GC content and presence of G4 sequences on the template strand around 3’-UTR regions (upstream of the TTS). The points represent genes, of which some contain G4 sequence(s) on the template strand upstream of the TTS (dark grey fill), while others do not (light grey fill). The x-axis indicates the GC percentage. The y-axis indicates the chance that a gene is differentially expressed between BS cells and controls as the negative natural logarithm of the raw p-values, with positive values indicating upregulation in BS cells, and negative values indicating downregulation in BS cells. An apparent trend is observed where genes with lower GC content are more frequently upregulated in BS fibroblasts.

Because different mechanisms could be responsible for up– and downregulation of transcription in BS cells, we analyzed the up– and downregulated gene sets separately as well (Supplementary Figures S2A and B). We also fitted regression models where we replaced the predictor G4 sequence presence by respectively the G4 sequences count or by a G4-score, calculated as the average mismatch percentage of each region (mismatch percentages served as the basis for identification of the experimentally defined G4 forming DNA sequences (13)) (Supplementary Figures S2C and D). The results of these additional analyses are in line with our findings described above.

### Transcriptome sequencing of human BS cells and Blm mutant mouse ES cells does not support a major role of local G4s in altering gene transcription in Blm mutants

Next, we examined newly generated RNA-seq profiles from cultured WT and BS lymphocytes and fibroblasts. Again, we used a linear model to estimate the relationship between the observed BS-specific transcriptional changes and the local presence of G4 sequences. For lymphocytes, this relationship was significant for G4 sequences on the template strand of the first intron and at either strand of the second and third intron and upstream and downstream of the TTS (Figure 2A). However, only G4 sequences on the coding strand upstream of the TTS remained significantly associated with the transcriptional alterations in BS when we controlled for GC content (Figure 2A). In fibroblasts, we found no significant effects of the presence of G4 sequences (Figure 2B) whereas we did find a significant effect of GC content upstream of the TTS. We also tested up– and downregulated genes separately for both cell types, and we repeated the regression analyses with alternative G4 definitions as well (Supplementary Figure 2) and found comparable results.

**Figure 2.**
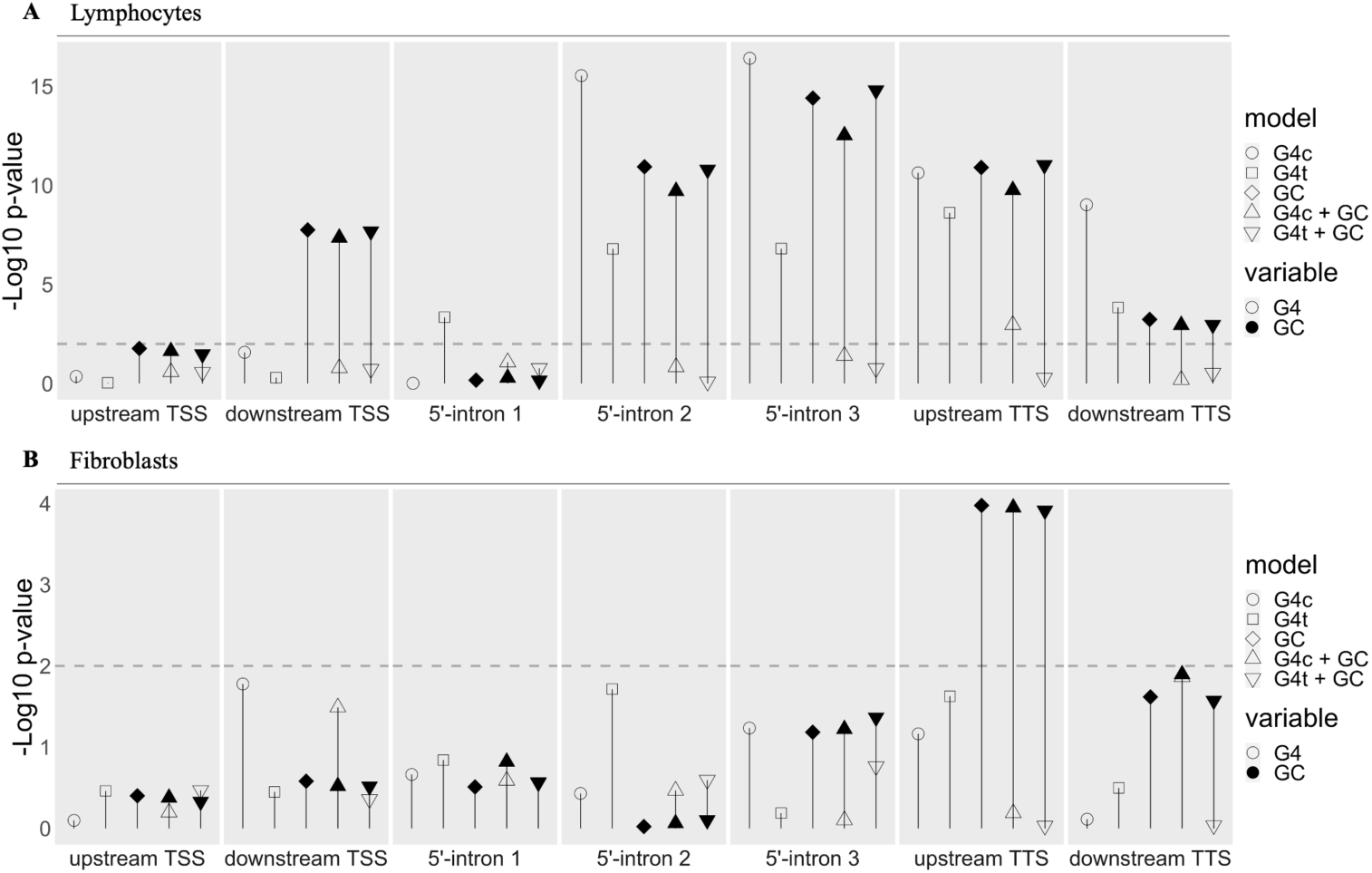
Results of regression analyses for BS-specific transcriptome alterations in RNA-seq data of lymphocytes and fibroblasts. (A). –log10 of the p-value for each variable, indicating the strength of the association of the presence of experimentally confirmed G4-forming sequences (no fill) on either the template (G4t) or coding strand (G4c) and GC content (black fill) in various gene regions with the transcriptome alterations between BS lymphocytes and controls predicted by the regression model. The shape of the data points indicates the regression model: the G4 presence on the coding strand of each gene region (G4c, circle), the G4 presence on the template strand of each gene region (G4t, square), the GC content of each gene region (GC, diamond), both the G4 presence on the coding strand and the GC content of each gene region (G4c + GC, triangle pointing up) or both the G4 presence on the template strand and the GC content of each gene region (G4t + GC, triangle pointing down). The dashed grey line marks p-value=0.01. (B). –log10 of the p-value for each variable, indicating the strength of the association of the presence of experimentally confirmed G4-forming sequences (no fill) on either the template (G4t) or coding strand (G4c) and GC content (black fill) in various gene regions with the transcriptome alterations between BS fibroblasts and controls predicted by the regression model.

Interestingly, we found significant associations of local GC content with expression alterations between fibroblast control samples (Supplementary Figure 3), suggesting that the observed GC-specific effect on transcription may be unrelated to BLM deficiency.

It is possible that the biological and genetic variability among different cell lines might obscure the G-quadruplex-related effects of BLM on gene expression. To exclude these effects, one should employ a model with a polymorphic and well-characterized genetic background where expression of BLM can be manipulated. We used *Blm* knockout and control ES cells from an F1 hybrid mouse embryonic stem (ES) cell line (cross between 129/Sv and Cast/EiJ inbred strains) to further investigate the role of local G4 sequences on transcription in the absence of *Blm*. Similarly to the previous datasets, these data do not show a specific effect linked to G4 sequences once GC content is taken into account (Figure S4).

### Analysis of allelic expression of genes discordant for G4 motifs in Blm mutant hybrid mice suggests that the GC content-dependent bias in gene expression likely originates from a technical source

A unique feature of the 129/Sv and Cast/EiJ cross is that its parental strains have very divergent genetic backgrounds that, on average, differ by 1 substitution per 114 bp. This diversity is an order of magnitude higher than seen in human genomes and exhibits a large amount of discordant G4 motifs that are present in either paternal or maternal chromosomes. We assessed the relationship between allele-specific expression (ASE) in respectively the *Blm* mutant cells and the control cells, and the presence of discordant G4 motifs between the two alleles (Figure 3 and Supplementary Figure 5). As genomic maps of experimentally validated G4 forming DNA sequences in 129/Sv and Cast/EiJ strains of laboratory mice are currently unavailable, this analysis remains limited to in silico predicted G4 motifs. Our scan for G4 motifs (G_3+_N_1– 7_G_3+_N_1−7_G_3+_N_1−7_G_3+_) polymorphic between 129/Sv and Cast/EiJ genetic backgrounds identified 1,484 strain-specific motifs associated with transcripts of 1,223 autosomal genes (Supplementary Table 1). Regression analysis showed that only discordant G4 motifs on the coding strand upstream of the TSS (p = 0.0015 and 0.0018) and the average GC content at the start of the first intron and upstream the TTS were mildly associated with ASE in the *Blm* mutant cells (Figure 3). In the control cells, we found that only discordant G4 motifs on the template strand at the start of the third intron and the average GC content upstream of the TTS were mildly but significantly associated with ASE (Supplementary Figure 5). It is worth noting that, despite their nominal significance, most of the aforementioned associations would not be considered significant after multiple testing correction and thus likely represent chance events. This lack of a robust relationship between discordant G4 motifs and ASE in *Blm* mutant cells, once more, does not support a major role of local G-quadruplexes in gene regulation in BLM-deficient cells. Furthermore, the limited effect of GC content in the absence of technical noise (compare Supplementary Figure 4A to Figure 3 and Supplementary Figure 5) suggests that the observed association between the GC content of genes and expression alterations between libraries is not a consequence of biological origin, but likely to stem from a technical artifact.

**Figure 3.**
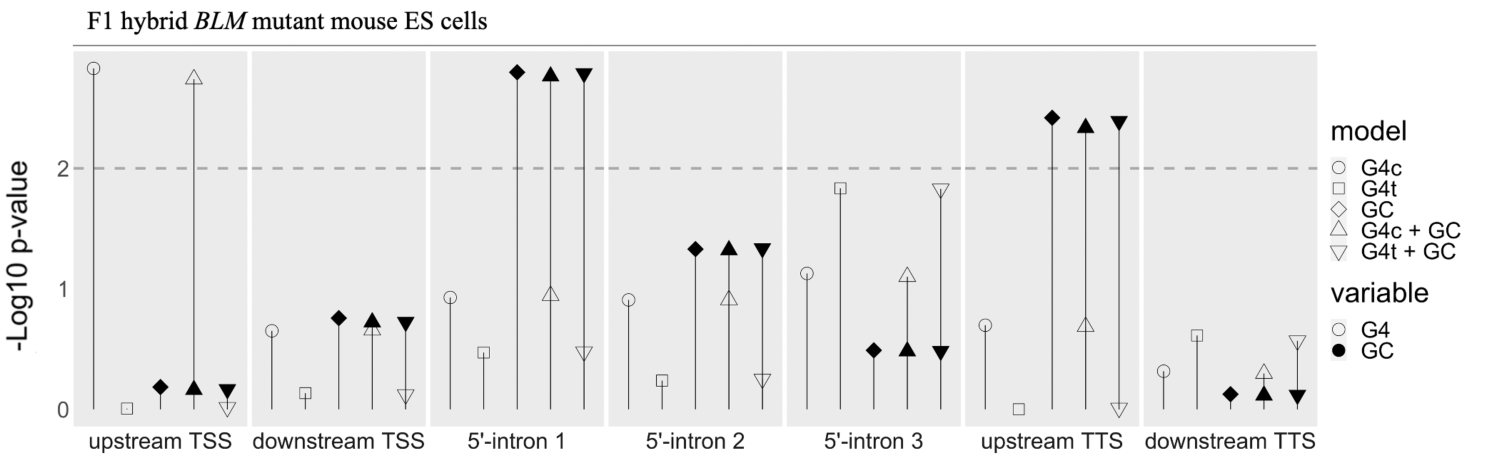
Results of regression analyses for allele-specific transcriptome alterations in RNA-seq data of F1 hybrid knockout mouse ES cells (129/Sv-Cast/EiJ). -log10 of the p-value for each variable, indicating the strength of the association of G4 motif discordance (–1, 0, or 1 when Cast/EiJ contained less, the same, or more G4 motifs than 129/Sv, no fill) on either the template (G4t) or coding strand (G4c) and GC content (average between 129/Sv and Cast/Eij strains, black fill) in various gene regions with allele-specific expression (ASE) in F1 hybrid mouse *Blm* knockout ES cell lines predicted by the regression model. The shape of the points indicates the regression model: the G4 motif discordance on the coding strand of each gene region (G4c, circle), the G4 motif discordance on the template strand on each gene region (G4t, square), the GC content of each gene region (GC, diamond), both the G4 motif discordance on the coding strand and the GC content of each gene region (G4c + GC, triangle pointing up) or both the G4 motif discordance on the template strand and the GC content of each gene region (G4t + GC, triangle pointing down). The dashed grey line marks p-value = 0.01.

### Allelic ratio at G4 motif sites discordant in F1 mice does not exhibit significant changes in expression level or variability

Another way to explore the effect of BLM deficiency at discordant G4 motifs is to compare allelic ratios between wild-type and *Blm* knockout cells. If alleles discordant for G4 motifs between CAST and 129 genetic backgrounds exhibit a common effect on expression in the absence of BLM, we should observe such an allelic difference between *Blm* knockout and WT mice (Figure 4A). Using ES cells from F1 mice, we checked whether the observed expression of the G4 motif-containing allele in the *Blm* knockout background differed from the expression that could be expected based on the G4 versus non-G4 alleles ratio seen in the wild-type cells. We did not observe any significant deviation of the observed-to-expected G4^+^_Blm KO_ ratios for G4 motifs discordant between 129 and CAST alleles (Figure 4B, C). The differences were insignificant for all gene partitions, at both coding and template strands, when testing means and when comparing variability to that of expressed genes where alleles are concordant for the G4 motif content.

**Figure 4.**
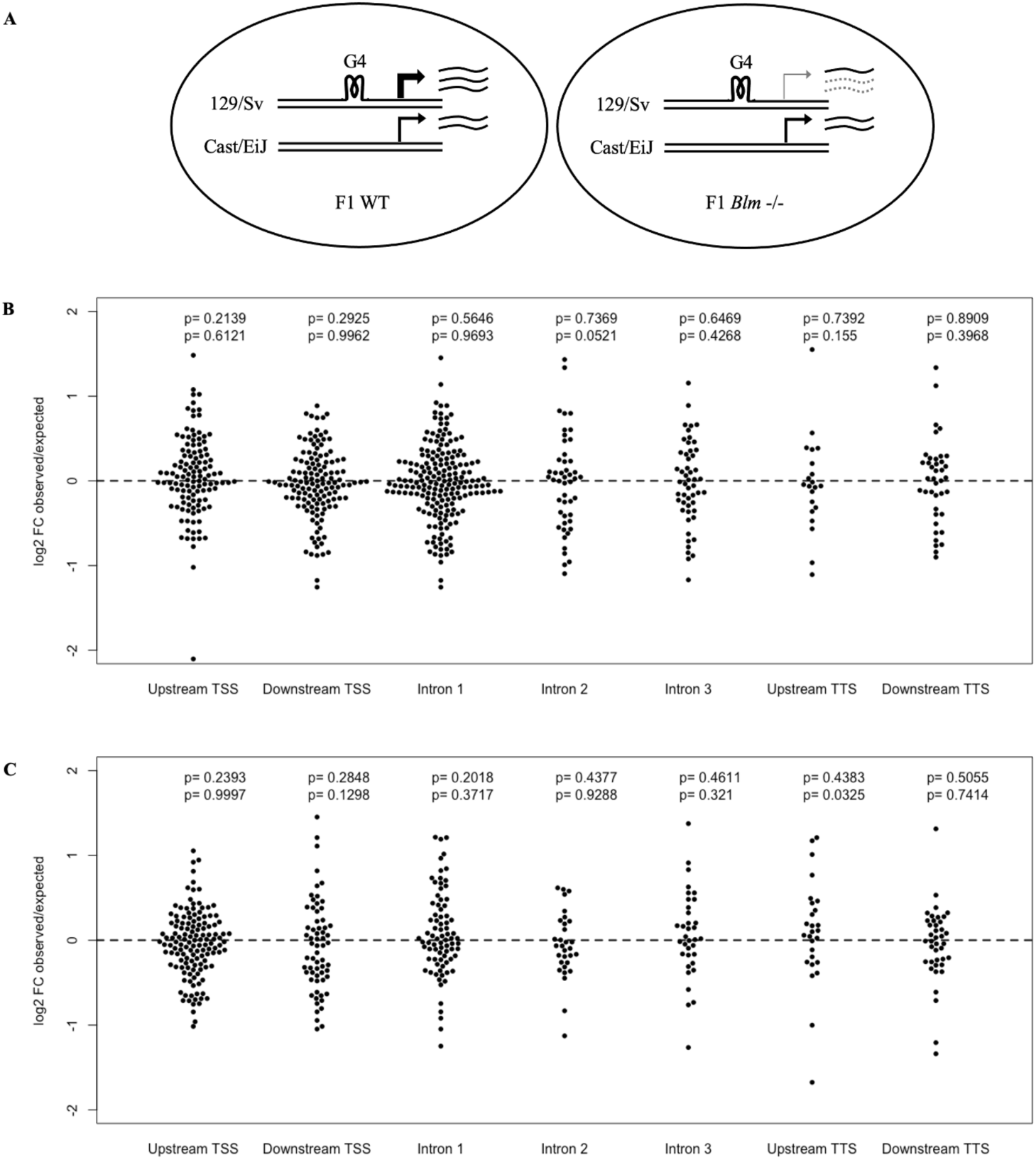
Analysis of gene expression in WT and *Blm* knockout cells in genes discordant for G4 motifs. (A) The BLM-specific effect that acts through G4 motifs can be observed as allelic disbalance in genes discordant for G4 motifs when allelic levels are compared between WT and *Blm* –/– samples. (B) Distribution of log_2_(observed/expected) expression ratio of G4 allele in *Blm* –/– cells when the G4 motif is present on coding strand. Nominal p-values correspond to the test of the mean (top row) or variance (bottom row) (C) Distribution of log_2_(observed/expected) expression ratio of G4 allele in *Blm* –/– cells when the G4 motif is present on the template strand. Nominal p-values correspond to the test of the mean (T-test, top row) or the variance (F-test, bottom row).

### BLM-deficient cells show an increase in exon skipping, with no evidence for enrichment of intronic G4 motifs

Apart from changing expression level of transcripts, G quadruplexes could potentially affect expression of transcript isoforms e.g., through regulation of alternative splicing. It was shown before that G4 motifs are overrepresented in 5’-intronic regions immediately flanking exons and thus could play a role in the regulation of splicing (15). We tested whether the absence of BLM has an effect on alternative splicing that is consistent among the datasets of our study. Although our analysis showed multiple hundreds of altered splicing events per dataset (Supplementary Table S2A), the vast majority of them were specific to a single dataset. Thus, we only found 13 events where the direction of the effect was the same in the human fibroblast and lymphocyte datasets (Supplementary Table S2B).

A common feature of all 3 datasets is a higher rate of exon exclusion in BLM-deficient samples (Supplementary Table S2A). In mutant samples, we observed more exon skipping events that reached FDR threshold of 0.01 compared to exon retention events (*X*^2^ test, p=2.18e-3; 6.68e-13; 2.96e-2 for fibroblast, lymphocyte, and ES-cell datasets respectively, Table S3). We, however, did not observe a robust relationship between the occurrence of exon skipping or exon retention events and the presence of G4 motifs in intronic flanks 250 bp downstream donor splice site of skipped, downstream or upstream exons (Supplementary Table S3).

### Clustering of similarly regulated genes suggests large-scale effects of BLM deficiency on gene expression

Several studies have suggested that the effect of G4s on gene expression can not only result from their direct effects on RNA polymerase processivity, but also through larger-scale mechanisms. Examples of such mechanisms could be the stabilization of nucleosome-depleted regions or increased stability of DNA loops that form topologically associated domains (16). These effects do not necessarily need to be confined to the body of the gene and can exert dysregulation of gene expression over longer genomic distances. We checked if genes up– or down-regulated in BLM-deficient samples tend to co-localize. Notably, differentially expressed gene neighbors frequently show the same direction of expression change (Table 1). The bias was not connected to whether genes are located on the same or opposite DNA strands (Supplementary Table S4).

**Table 1.** Analysis of change directionality of differentially expressed gene pairs, located within 100 kbp from each other.

| Dataset | Regulated in the same direction | Regulated in the opposite direction | P-value |
| --- | --- | --- | --- |
| Human fibroblasts | 1036 | 528 | $5.87 \cdot 10^{-20}$ |
| Human lymphocytes | 127 | 8 | $1.99 \cdot 10^{-10}$ |
| Mouse ES cells | 35 | 7 | $1.33 \cdot 10^{-3}$ |

Such a co-regulation gene expression pattern extends far beyond the size of a typical gene length (e.g. in mouse ES cells, Figure 5A, top panel). We observe large genomic segments where differentially regulated genes show a similar directionality of expression change. This observation suggests that BLM deficiency manifests itself at a larger, potentially (sub-) chromosomal scale that involves multiple genes, rather than being guided by G4 presence around or inside a gene body.

**Figure 5.**
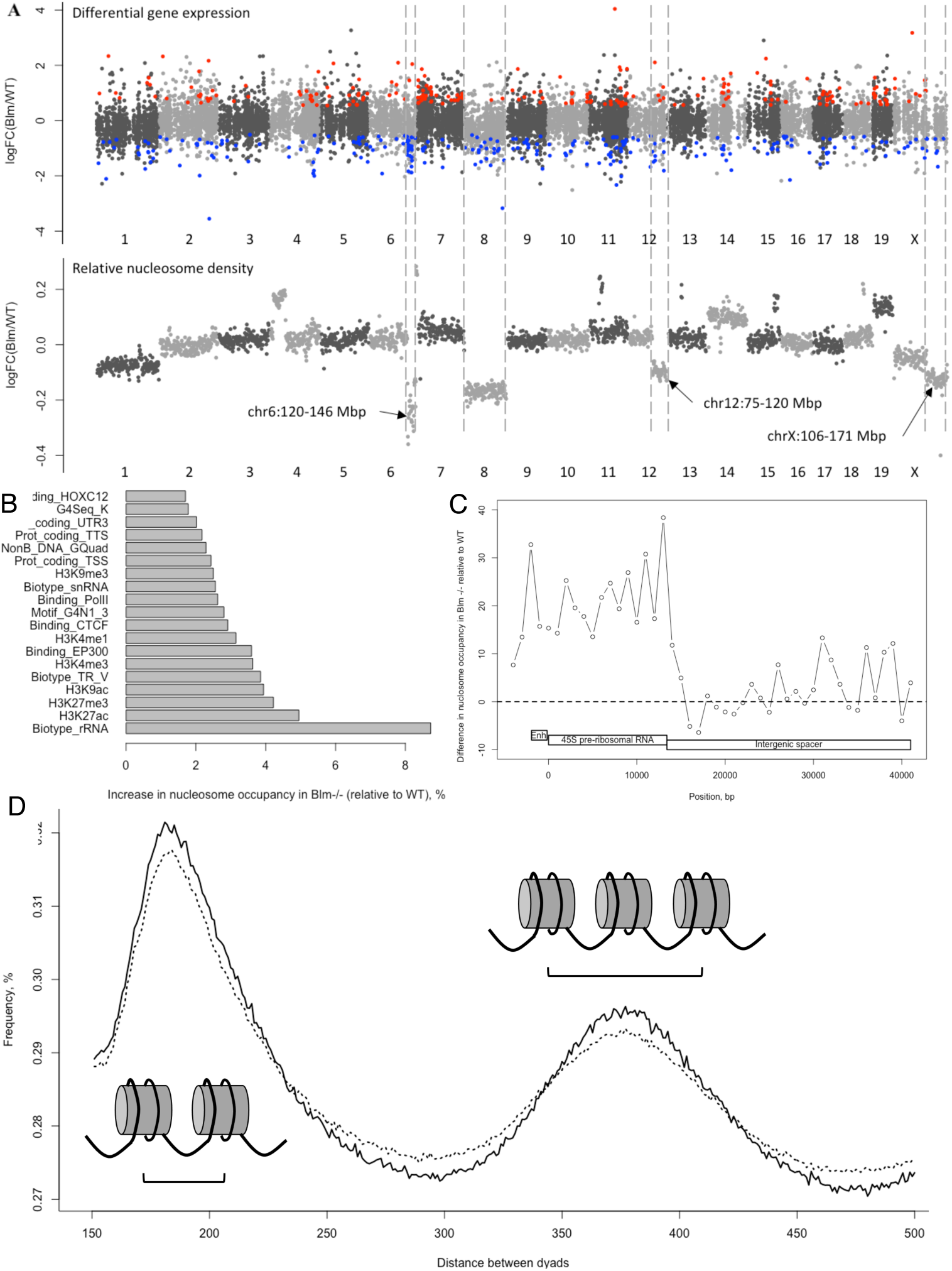
The large-scale structure of changes in gene expression and nucleosome occupancy in *Blm* knockout cells. (A) Significantly up– and down-regulated genes in *Blm* –/– cells (given by red and blue circles respectively) are clustered in relation to their chromosomal location (top panel) and relative nucleosome density (bottom panel). Horizontal positions show normalized relative log_2_ fold change per gene (expression, top panel) or log_2_ fold change in nucleosome occupancy per 1 megabase window (bottom panel). (B) Top 20 genomic features showing increased nucleosome occupancy in Blm –/-, compared to wild-type mouse ES cells (C) Local nucleosome density in *Blm* –/– ES cells (per 1000 bp, percent difference relative to WT cells) in 45kb mouse rRNA repeat (GenBank acc. BK000964) (D) Frequency distribution of the relative distance between nucleosome centers for distance range 150 – 500 bp. Higher peaks and deeper valleys for wild-type cells (solid line) reflect stricter nucleosome phasing compared to that of *Blm* knockout cells (dotted line).

### Nucleosome occupancy analysis shows higher density at the pre-ribosomal RNA gene and less phasing in F1 Blm mutants

The observed long-range unidirectional transcription changes might involve large-scale alterations in chromatin structure. We used Strand-seq data generated for the same ES cells (7) to investigate differences in nucleosome occupancy between *Blm-/-* and WT samples. We observe segmental changes spanning megabase-long DNA segments (Figure 5A, bottom panel) that indicate global changes in chromatin structure rather than by presence of local G-quaruplexes.

In order to connect changes in chromatin packaging with genomic features, we checked whether nucleosomes are depleted at G4 sites in *Blm* knockout cells, which would be expected if G4 structures are stabilized. We discarded the hypothesis as nucleosomes had slightly more preference for putative G4 sequences in *Blm*-/– (6.86% of reads) rather than wild-type cells (6.83% of reads). Further, we observed little difference in other parameters for nucleosome-bound regions in *Blm-/-* and WT cells: GC content (average 42.35% and 42.36% respectively) or overlap of CpG islands (0.399% vs 0.396% of reads). We also tested differential occupancy for gene bodies, exons, or other partitions that we considered above, but did not observe significant differences between nucleosome profiles of wildtype and knockout samples (Supplementary Table S5). Finally, we explored if gene biotype, epigenetic modifications, binding sites for transcriptional factors, and non-B DNA regions exhibited differential nucleosome occupancy. Interestingly, some of these features showed a higher nucleosome density in mutant cells (Figure 5B, Supplementary Table S6). The greatest difference (+8.72% in *Blm*-/-) was observed in rRNA gene bodies followed by several types of epigenetic marks: H3K27ac (+4.95%, associated with active enhancer regions), H3K27me3 (+4.22%, linked to heterochromatin formation), H3K9ac (+3.94%, connected with active promotors). To zoom in on the rRNA locus (this highly repetitive cluster is poorly represented in mouse genome reference), we remapped our nucleosome footprint data to a genome reference containing a consensus rRNA repeat unit. We observed about 20% increase in nucleosome occupancy at upstream regulatory and 45S pre-ribosomal RNA in *Blm* mutant cells that does not seem to extend to the intergenic spacer (Figure 5B).

The higher nucleosome density at the rRNA cluster in *Blm* mutants could result in repressed transcription of rRNA and, as a result, have a negative effect on global translation and systemically alter gene regulation in *Blm*-/– cells.

Lastly, we questioned if nucleosome phasing is globally different between mutant and wild-type cells. The perturbation of nucleosome positioning is linked to changes in gene expression, where active genes typically show more strict phasing. We observe that *Blm* knockout cells show deterioration of nucleosome phasing (Figure 5D) with overall less predictable relative nucleosome positioning compared to wild-type cells.

In conclusion, our analysis of a broad set of genomic features suggests that pre-ribosomal RNA and not G-quadruplexes act as the strongest effector of BLM deficiency. The resulting changes in chromatin and gene expression span large chromosomal regions and are coordinated whereby the presence of individual G4s in a gene body is unlikely to play a major role.

## Discussion

The G4-resolving abilities of the BLM helicase have led to the implication of dysregulated G4 formation as a potential causal factor for the Bloom Syndrome phenotype (7, 8, 10, 11, 17). One process through which unresolved G4s have been proposed to inflict the disease is transcription (10, 11), as BS-specific transcriptional differences were reportedly correlated with the presence of G4 sequences in certain gene regions. The studies that reported these correlations made use of *in silico* predicted G4 locations (10, 11), as the number of experimentally observed G4 sequences used to be very limited. However, this set was greatly expanded recently (12, 13), enabling us to comprehensively investigate the relationship between the local presence of these experimentally defined G4 sequences and BS-specific transcriptional alterations. Unlike previous studies, we have chosen to employ linear regression analysis to explicitly correct for local GC content and to avoid the use of arbitrary cut-offs for differential expression.

While a previous study concluded that BLM helicase regulates gene expression and that one process by which this regulation occurs may involve direct binding to G-quadruplex structures (11), the results of our reanalysis of public data and additionally generated transcriptome data from both BS patients and murine *Blm* mutant cells do not agree with this hypothesis. Our conclusion is based on an analysis of several datasets encompassing different experimental platforms and species. We reanalyzed a previously reported microarray cohort of fourteen BS fibroblast cell lines and twelve controls (11), and found that the effect of local G4 sequences on BS-specific transcription disappeared completely when we account for differences in GC content. We also profiled the transcriptomes from EBV-transformed B-lymphocyte and primary fibroblast BS lines using transcriptome sequencing but did not find strong evidence for a role of sequences with the potential to form G4s in altering gene expression in the absence of BLM. Furthermore, the assessment of the role of G4 sequences in the direct effect of *Blm* knockout on transcription in an F1 hybrid mouse ES cell line (129Sv-Cast/EiJ) did not provide any support for the involvement of local G-quadruplexes in gene regulation in BLM deficient cells.

Regarding the role of GC content, we found that the local GC percentage significantly correlated with transcriptional differences not only between BLM deficient and wild-type samples but also when control samples are compared. This suggests that the observed GC effect on transcription may be unrelated to BLM deficiency. The existence of such a sample-specific GC effect in RNA-seq data has been observed before by several researchers and has mostly been regarded as a technical artifact (18, 19, 20). And indeed, we only observed a nominally significant association between GC content and allele-specific expression (ASE) in the F1 hybrid *Blm* mutant and wildtype cells. Since the allele-specific measurements can be considered as free from the technical variability among experiments, our results indicate that the GC effect is likely to be a consequence of a technical bias rather than a result of the absence of functional BLM helicase.

Although likely unrelated to local G-quadruplexes, our analysis does identify substantial differences between BLM-deficient and WT transcriptomes (Supplementary Figure 6). Genes upregulated in BLM-deficient cells share similar expression patterns between cell types (fibroblasts, lymphocytes) and platforms (expression arrays, RNA-seq). Similarly, our analysis of alternative splicing identifies exon skipping as a type of event, overrepresented in Blm mutant compared to wild-type cells. A matching trend has been observed in a recent study using human neuronal cells that show a significant overrepresentation of G4-containing introns among exon-skipping events after depolarizing KCl treatment (that can stabilize G4 structures) (21). Our tests, however, do not show significant enrichment of intronic G4s flanking either the skipped, previous or next exon of the same transcript. Based on the observation of common regulatory changes in the level and structure of transcripts seen in BLM deficient cells, and the absence of a strong correlation to G4 presence, our results suggest that DNA quadruplex structures are not the primary mediators of expression changes in *Blm –/-* cells.

Another strong argument against the major role of gene-specific G4 sites is a long-range span of regulatory changes seen between *Blm-/-* and WT cells. Differentially expressed gene neighbors frequently showed the same direction of expression change and this pattern was observed in all three RNA-seq datasets. This concerted change is supported by an analysis of nucleosome density showing stable levels of nucleosome density across large genomic segments. To investigate what might be responsible for the observed large-scale changes in chromatin structure we surveyed various types of genomic features – GC content, CpG islands, different non-B DNA regions (including G4s), epigenetic modifications, transcription factor binding sites, and gene features and biotypes. Interestingly, the genomic feature that shows the strongest effect in the *Blm*-/– cells was the ribosomal RNA locus. Our fine-mapping analysis showed that a denser histone packaging is characteristic for the 45S pre-ribosomal RNA unit of the locus.

The connection between BLM and rRNA was observed in multiple studies. Thus, *in situ* hybridizations showed that, during the S phase, BLM can be found in the nucleolus (22) and that its presence facilitates rRNA transcription (23). Further, while previous research showed that, within the rDNA cluster, BLM localizes primarily to 18S and Alu repeat regions (24), we observe that the change in chromatin packaging specific for the BLM mutant encompasses the pre-rRNA region and its upstream region.

Recent studies of 3D chromatin organization indicate that the rDNA locus can play an important role in global regulation of gene expression during development, e.g. through contacts to H3K27ac-marked super-enhancers (reviewed by (25)). In support of this hypothesis, we observed H3K27ac regions as the second most strongly affected genomic feature, having a 5% increase in nucleosome density in the absence of BLM. We, therefore, suggest that future studies should explore the role of global chromatin organization in Bloom syndrome.

## Data availability

RNA-seq data reported in this paper have been submitted to the ArrayExpress database (http://www.ebi.ac.uk/arrayexpress) under accession number E-MTAB-11131. The code generated during this study is available at GitHub; https://github.com/thamarlobo/BLM_analysis.

## Author contributions

TJL and VG designed the study. TJL and VG performed data analysis. TJL wrote the paper. PML and VG edited the manuscript. PML and VG jointly supervised the project.

## Acknowledgments

We thank Dr. N. van Wietmarschen for generating the sequencing libraries, and we thank Dr. D.C.J. Spierings and N. Halsema and other members of ERIBA sequencing facility for their help with sequencing the samples. We are also grateful to Dr. M. Schubert and Dr. YM. Moshkin for useful discussions.

## Supporting information

### Supplementary Figures

**Figure S1.**
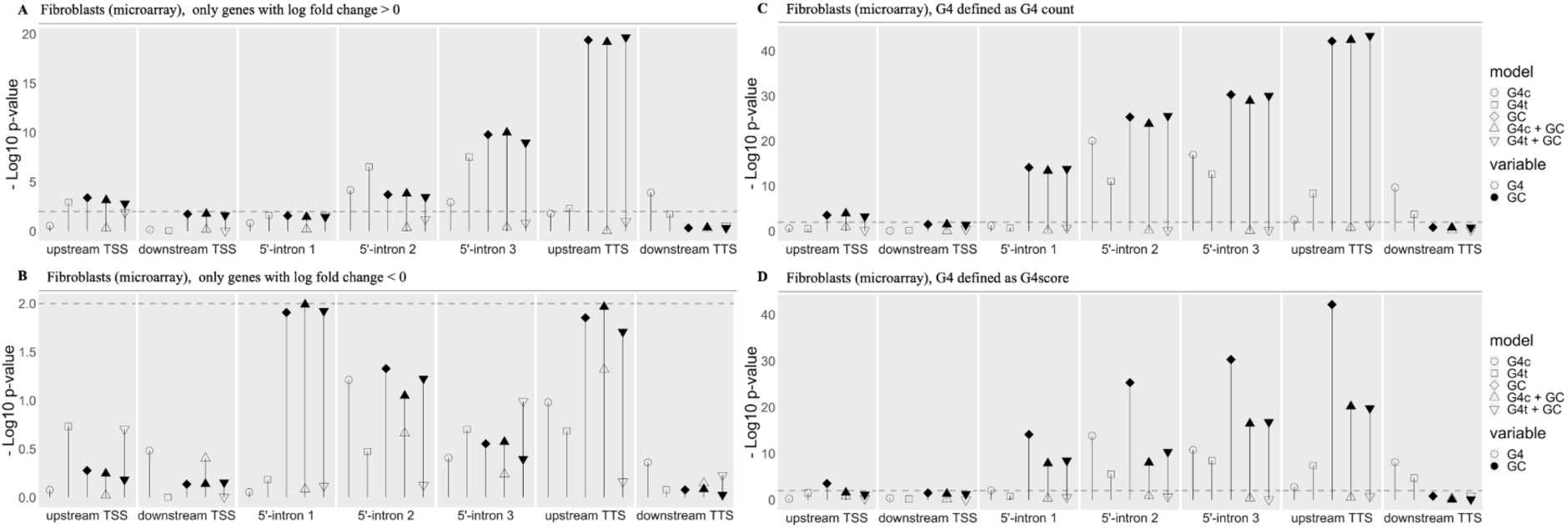
Results of regression analyses for BS-specific transcriptome alterations in microarray data of fibroblasts. (A). – log10 of the p-value for each variable, indicating the strength of the association of the presence of experimentally confirmed G4-forming sequences (no fill) on either the template (G4t) or coding strand (G4c) and GC content (black fill) in various gene regions with the upregulation of genes in BS fibroblasts predicted by the regression model. The shape of the points indicates the regression model: the G4 presence on the coding strand of each gene region (G4c, circle), the G4 presence on the template strand on each gene region (G4t, square), both the G4 presence on the coding strand and the GC content of each gene region (G4c + GC, triangle point up) or both the G4 presence on the template strand and the GC content of each gene region (G4t + GC, triangle point down). The dashed grey line marks p-value = 0.01. (B). –log10 of the p-value for each variable, indicating the strength of the association of the presence of experimentally confirmed G4-forming sequences (no fill) on either the template (G4t) or coding strand (G4c) and GC content (black fill) in various gene regions with the downregulation of genes in BS fibroblasts, predicted by the regression model. The shape of the points indicates the regression model: the G4 presence on the coding strand of each gene region (G4c, circle), the G4 presence on the template strand on each gene region (G4t, square), both the G4 presence on the coding strand and the GC content of each gene region (G4c + GC, triangle point up) or both the G4 presence on the template strand and the GC content of each gene region (G4t + GC, triangle point down). The dashed grey line marks p-value = 0.01. (C). –log10 of the p-value for each variable, indicating the strength of the association of experimentally confirmed G4-forming sequence count (no fill) on either the template (G4t) or coding strand (G4c) and GC content (black fill) in various gene regions with the transcriptome alterations between BS fibroblasts and controls predicted by the regression model. The shape of the points indicates the regression model: the G4 count on the coding strand of each gene region (G4c, circle), the G4 count on the template strand on each gene region (G4t, square), both the G4 count on the coding strand and the GC content of each gene region (G4c + GC, triangle point up) or both the G4 count on the template strand and the GC content of each gene region (G4t + GC, triangle point down). The dashed grey line marks p-value = 0.01. (D). –log10 of the p-value for each variable, indicating the strength of the association of the G4score (the average mismatch percentage as determined by Marsico et al., 2019) (no fill) on either the template (G4t) or coding strand (G4c) and GC content (black fill) in core various gene regions with transcriptome alterations between BS fibroblasts and controls predicted by the regression model. The shape of the points indicates the regression model: the G4score on the coding strand of each gene region (G4c, circle), the G4score on the template strand on each gene region (G4t, square), both the G4score on the coding strand and the GC content of each gene region (G4c + GC, triangle point up) or both the G4score on the template strand and the GC content of each gene region (G4t + GC, triangle point down). The dashed grey line marks p-value = 0.01.

**Figure S2.**
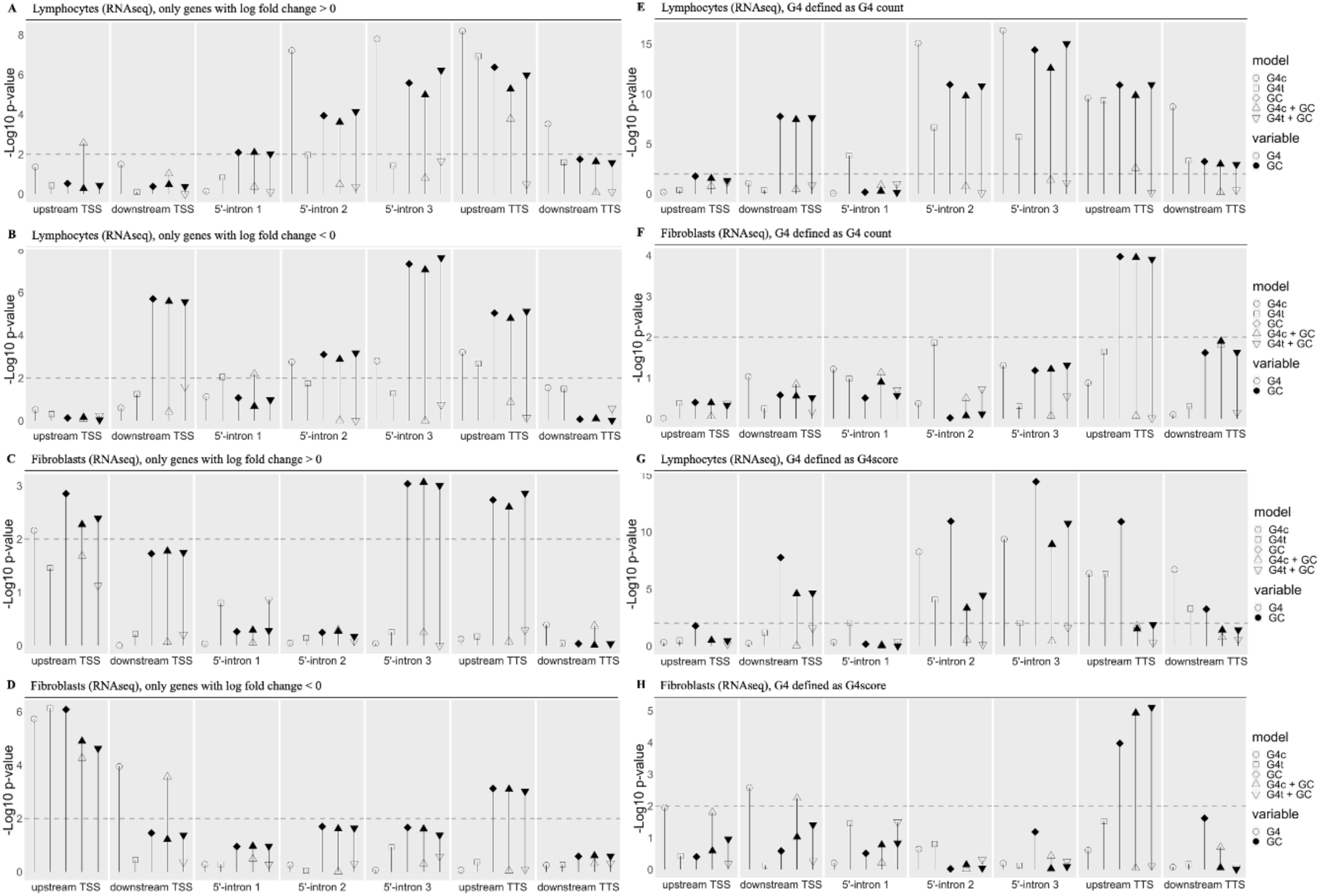
Results of regression analyses for BS-specific transcriptome alterations in RNA-seq data of lymphocytes and fibroblasts. (A). –log10 of the p-value for each variable, indicating the strength of the association of the presence of experimentally confirmed G4-forming sequences (no fill) on either the template (G4t) or coding strand (G4c) and GC content (black fill) in various gene regions with the upregulation of genes in BS lymphocytes predicted by the regression model. The shape of the points indicates the regression model: the G4 presence on the coding strand of each gene region (G4c, circle), the G4 presence on the template strand on each gene region (G4t, square), both the G4 presence on the coding strand and the GC content of each gene region (G4c + GC, triangle point up) or both the G4 presence on the template strand and the GC content of each gene region (G4t + GC, triangle point down). The dashed grey line marks p-value = 0.01. (B). –log10 of the p-value for each variable, indicating the strength of the association of the presence of experimentally confirmed G4-forming sequences (no fill) on either the template (G4t) or coding strand (G4c) and GC content (black fill) in various gene regions with the downregulation of genes in BS lymphocytes predicted by the regression model. The shape of the points indicates the regression model: the G4 presence on the coding strand of each gene region (G4c, circle), the G4 presence on the template strand on each gene region (G4t, square), both the G4 presence on the coding strand and the GC content of each gene region (G4c + GC, triangle point up) or both the G4 presence on the template strand and the GC content of each gene region (G4t + GC, triangle point down). The dashed grey line marks p-value = 0.01. (C). –log10 of the p-value for each variable, indicating the strength of the association of the presence of experimentally confirmed G4-forming sequences (no fill) on either the template (G4t) or coding strand (G4c) and GC content (black fill) in various gene regions with the upregulation of genes in BS fibroblasts predicted by the regression model. The shape of the points indicates the regression model: the G4 presence on the coding strand of each gene region (G4c, circle), the G4 presence on the template strand on each gene region (G4t, square), both the G4 presence on the coding strand and the GC content of each gene region (G4c + GC, triangle point up) or both the G4 presence on the template strand and the GC content of each gene region (G4t + GC, triangle point down). The dashed grey line marks p-value = 0.01. (D). –log10 of the p-value for each variable, indicating the strength of the association of the presence of experimentally confirmed G4-forming sequences (no fill) on either the template (G4t) or coding strand (G4c) and GC content (black fill) in various gene regions with the downregulation of genes in BS fibroblasts predicted by the regression model. The shape of the points indicates the regression model: the G4 presence on the coding strand of each gene region (G4c, circle), the G4 presence on the template strand on each gene region (G4t, square), both the G4 presence on the coding strand and the GC content of each gene region (G4c + GC, triangle point up) or both the G4 presence on the template strand and the GC content of each gene region (G4t + GC, triangle point down). The dashed grey line marks p-value = 0.01. (E). –log10 of the p-value for each variable, indicating the strength of the association of experimentally confirmed G4-forming sequence count (no fill) on either the template (G4t) or coding strand (G4c) and GC content (black fill) in various gene regions with the transcriptome alterations between BS lymphocytes and controls predicted by the regression model. The shape of the points indicates the regression model: the G4 count on the coding strand of each gene region (G4c, circle), the G4 count on the template strand on each gene region (G4t, square), both the G4 count on the coding strand and the GC content of each gene region (G4c + GC, triangle point up) or both the G4 count on the template strand and the GC content of each gene region (G4t + GC, triangle point down). The dashed grey line marks p-value = 0.01. (F). –log10 of the p-value for each variable, indicating the strength of the association of experimentally confirmed G4-forming sequence count (no fill) on either the template (G4t) or coding strand (G4c) and GC content (black fill) in various gene regions with the transcriptome alterations between BS fibroblasts and controls predicted by the regression model. The shape of the points indicates the regression model: the G4 count on the coding strand of each gene region (G4c, circle), the G4 count on the template strand on each gene region (G4t, square), both the G4 count on the coding strand and the GC content of each gene region (G4c + GC, triangle point up) or both the G4 count on the template strand and the GC content of each gene region (G4t + GC, triangle point down). The dashed grey line marks p-value = 0.01. (G). –log10 of the p-value for each variable, indicating the strength of the association of G4score (the average mismatch percentage as determined by Marsico et al., 2019) (no fill) on either the template (G4t) or coding strand (G4c) and GC content (black fill) in various gene regions with the transcriptome alterations between BS lymphocytes and controls predicted by the regression model. The shape of the points indicates the regression model: the G4score on the coding strand of each gene region (G4c, circle), the G4score on the template strand on each gene region (G4t, square), both the G4score on the coding strand and the GC content of each gene region (G4c + GC, triangle point up) or both the G4score on the template strand and the GC content of each gene region (G4t + GC, triangle point down). The dashed grey line marks p-value = 0.01. (H). –log10 of the p-value for each variable, indicating the strength of the association of G4score (the average mismatch percentage as determined by Marsico et al., 2019) (no fill) on either the template (G4t) or coding strand (G4c) and GC content (black fill) in various gene regions with the transcriptome alterations between BS fibroblasts and controls predicted by the regression model. The shape of the points indicates the regression model: the G4score on the coding strand of each gene region (G4c, circle), the G4score on the template strand on each gene region (G4t, square), both the G4score on the coding strand and the GC content of each gene region (G4c + GC, triangle point up) or both the G4score on the template strand and the GC content of each gene region (G4t + GC, triangle point down). The dashed grey line marks p-value = 0.01.

**Figure S3.**
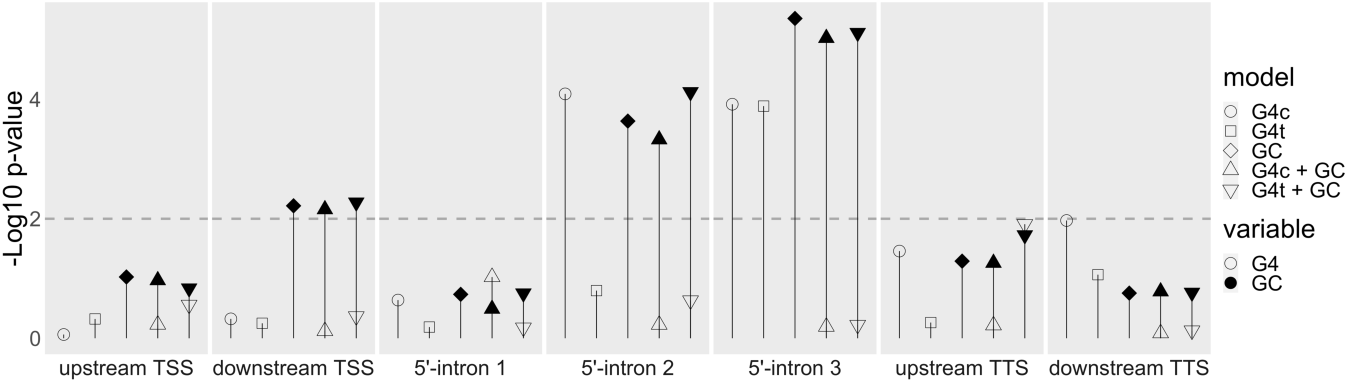
Results of regression analyses for transcriptome alterations between control cell lines in RNA-seq data of fibroblasts. -log10 of the p-value for each variable, indicating the strength of the association of the presence of G4 motifs (no fill) on either the template (G4t) or coding strand (G4c) and GC content (black fill) in various gene regions with the transcriptome alterations between fibroblast control cell lines predicted by the regression model. The shape of the points indicates the regression model: the presence of G4 motifs on the coding strand of each gene region (G4c, circle), the presence of G4 motifs on the template strand on each gene region (G4t, square), the GC content of each gene region (GC, diamond), both the presence of G4 motifs on the coding strand and the GC content of each gene region (G4c + GC, triangle point up) or both the presence of G4 motifs on the template strand and the GC content of each gene region (G4t + GC, triangle point down). The dashed grey line marks p-value=0.01.

**Figure S4.**
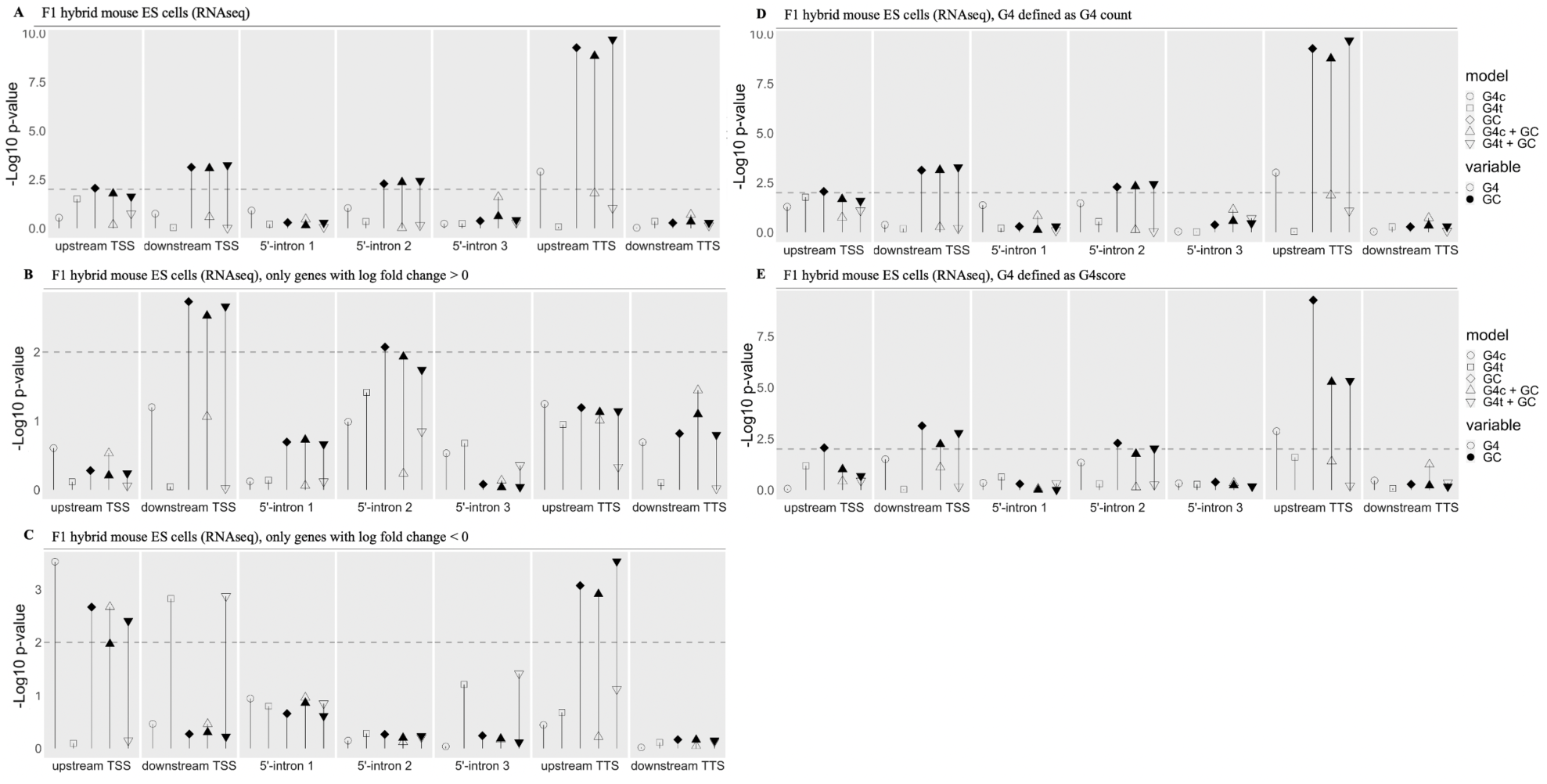
Results of regression analyses for *Blm* knockout-specific transcriptome alterations in RNA-seq data of F1 hybrid mouse ES cells (129Sv-Cast/EiJ). (A). – log10 of the p-value for each variable, indicating the strength of the association of the presence of experimentally confirmed G4-forming sequences (no fill) on either the template (G4t) or coding strand (G4c) and GC content (black fill) in various gene regions with the transcriptome alterations between F1 hybrid mouse *Blm* knockout ES cell lines and controls predicted by the regression model. The shape of the points indicates the regression model: the G4 presence of G4 motifs on the coding strand of each gene region (G4c, circle), the G4 presence of G4 motifs on the template strand of each gene region (G4t, square), the GC content of each gene region (GC, diamond), both the G4 presence of G4 motifs on the coding strand and the GC content of each gene region (G4c + GC, triangle pointing up) or both the G4 presence on the template strand and the GC content of each gene region (G4t + GC, triangle pointing down). The dashed grey line marks p-value=0.01. (B). –log10 of the p-value for each variable, indicating the strength of the association of the presence of experimentally confirmed G4-forming sequences (no fill) on either the template (G4t) or coding strand (G4c) and GC content (black fill) in various gene regions with the upregulation of genes in F1 hybrid mouse *Blm* knockout ES cell lines predicted by the regression model. The shape of the points indicates the regression model: the G4 presence on the coding strand of each gene region (G4c, circle), the G4 presence on the template strand on each gene region (G4t, square), both the G4 presence on the coding strand and the GC content of each gene region (G4c + GC, triangle point up) or both the G4 presence on the template strand and the GC content of each gene region (G4t + GC, triangle point down). The dashed grey line marks p-value = 0.01. (C). –log10 of the p-value for each variable, indicating the strength of the association of the presence of experimentally confirmed G4-forming sequences (no fill) on either the template (G4t) or coding strand (G4c) and GC content (black fill) in various gene regions with the downregulation of genes in F1 hybrid mouse *Blm* knockout ES cell lines predicted by the regression model. The shape of the points indicates the regression model: the G4 presence on the coding strand of each gene region (G4c, circle), the G4 presence on the template strand on each gene region (G4t, square), both the G4 presence on the coding strand and the GC content of each gene region (G4c + GC, triangle point up) or both the G4 presence on the template strand and the GC content of each gene region (G4t + GC, triangle point down). The dashed grey line marks p-value = 0.01. (D). –log10 of the p-value for each variable, indicating the strength of the association of G4 motif count (no fill) on either the template (G4t) or coding strand (G4c) and GC content (black fill) in various gene regions with the transcriptome alterations between F1 hybrid mouse *Blm* knockout ES cell lines and controls predicted by the regression model. The shape of the points indicates the regression model: the G4 motif count on the coding strand of each gene region (G4c, circle), the G4 motif count on the template strand on each gene region (G4t, square), both the G4 motif count on the coding strand and the GC content of each gene region (G4c + GC, triangle point up) or both the G4 motif count on the template strand and the GC content of each gene region (G4t + GC, triangle point down). The dashed grey line marks p-value = 0.01. (E). –log10 of the p-value for each variable, indicating the strength of the association of the G4score (the average mismatch percentage as determined by Marsico et al., 2019) (no fill) on either the template (G4t) or coding strand (G4c) and GC content (black fill) in core various gene regions with transcriptome alterations between F1 hybrid mouse *Blm* knockout ES cell lines and controls predicted by the regression model. The shape of the points indicates the regression model: the G4score on the coding strand of each gene region (G4c, circle), the G4score on the template strand on each gene region (G4t, square), both the G4score on the coding strand and the GC content of each gene region (G4c + GC, triangle point up) or both the G4score on the template strand and the GC content of each gene region (G4t + GC, triangle point down). The dashed grey line marks p-value = 0.01.

**Figure S5.**
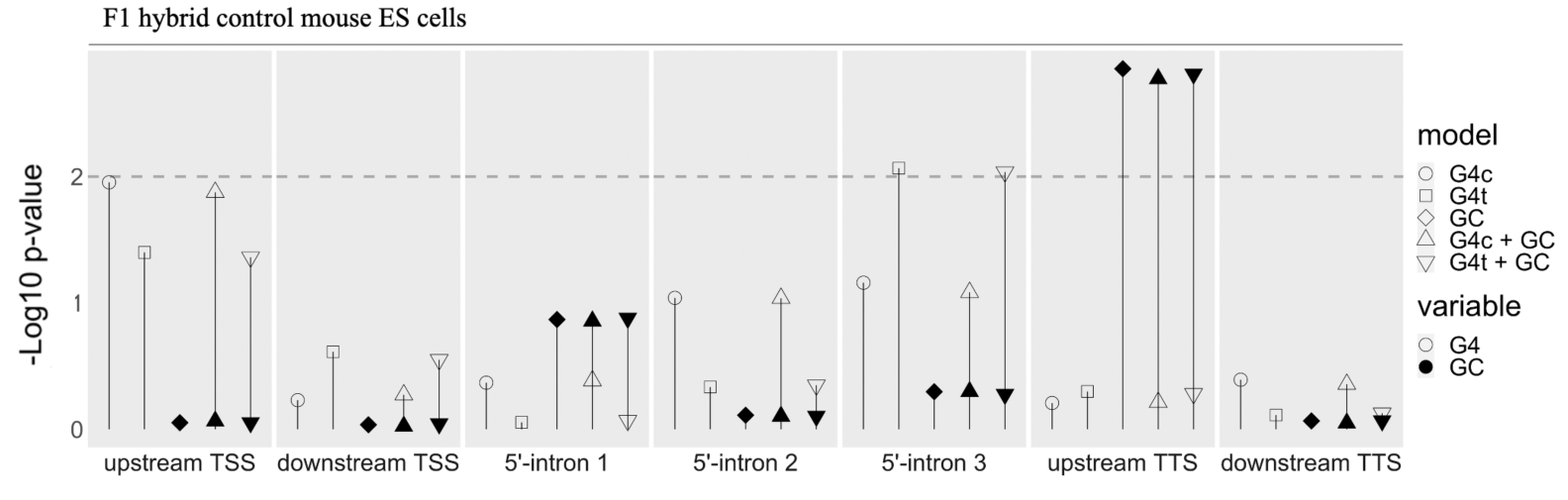
Results of regression analyses for allele-specific transcriptome alterations in RNA-seq data of F1 hybrid knockout mouse ES cells (129/Sv-Cast/EiJ). -log10 of the p-value for each variable, indicating the strength of the association of G4 motif discordance (–1, 0, or 1 when Cast/EiJ contained less, the same, or more G4 motifs than 129/Sv, no fill) on either the template (G4t) or coding strand (G4c) and GC content (average between 129/Sv and Cast/Eij strains, black fill) in various gene regions with allele-specific expression (ASE) in F1 hybrid mouse control ES cell lines predicted by the regression model. The shape of the points indicates the regression model: the G4 motif discordance on the coding strand of each gene region (G4c, circle), the G4 motif discordance on the template strand on each gene region (G4t, square), the GC content of each gene region (GC, diamond), both the G4 motif discordance on the coding strand and the GC content of each gene region (G4c + GC, triangle pointing up) or both the G4 motif discordance on the template strand and the GC content of each gene region (G4t + GC, triangle pointing down). The dashed grey line marks p-value = 0.01.

**Figure S6.**
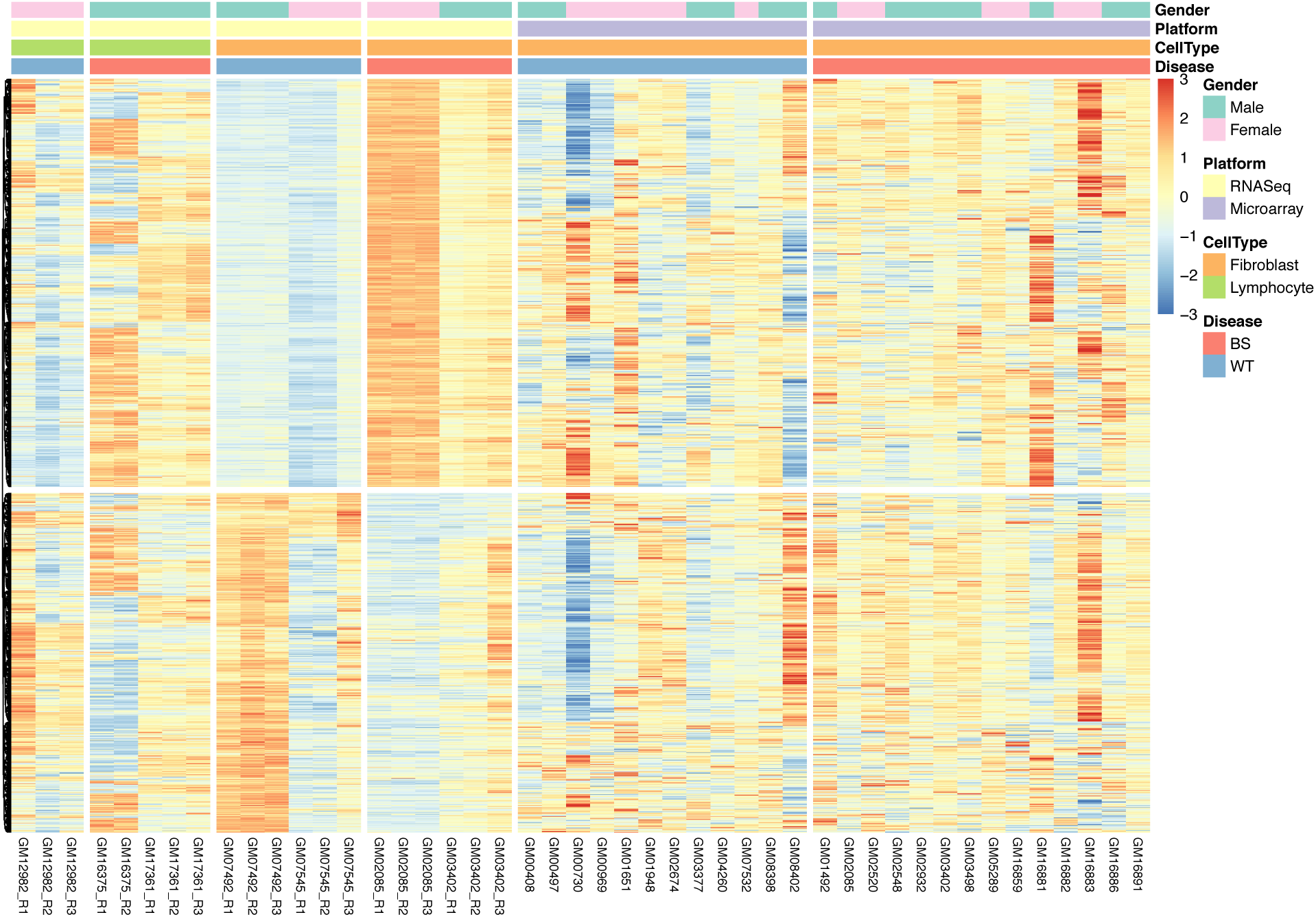
Heatmap of genes significant between BS and WT samples. Genes with FDR <=0.01 were selected from 3 experiments on human cells. Z-scores based on normalized expression values were used. Columns represent experimental conditions. Upper blocks significantly upregulated in BLM deficient cells and lower block – downregulated genes.

### Supplementary Tables

**Supplementary Table S1.** Numbers of G4 motifs (G_3+_N_1–7_G_3+_N_1−7_G_3+_N_1−7_G_3+_) identified as polymorphic between 129 and CAST genetic backgrounds. We scanned genomes of 129/Sv and Cast/Eij strains of laboratory mice for discordant G4 motifs (G_3+_N_1–7_G_3+_N_1−7_G_3+_N_1−7_G_3+_) and found the following numbers of autosomal genes with discordant G4 motifs in the investigated regions.

| Region | Total number of genes with discordant G4 motifs in this region | Total number of discordant G4 motifs in this region |
| --- | --- | --- |
| upstream TSS | 335 | 338 |
| downstream TSS | 277 | 282 |
| 5'-intron 1 | 355 | 362 |
| 5'-intron 2 | 130 | 167 |
| 5'-intron 3 | 117 | 119 |
| upstream TTS | 70 | 70 |
| downstream TTS | 145 | 146 |

**Supplementary Table S2A.** Summary of RMATS analysis of alternative splicing. Number of significant events (FDR<0.01) per dataset, event type and direction.

| Type of event | Observation | Human fibroblasts |  | Human lymphocytes |  | Mouse ES cells |  |
| --- | --- | --- | --- | --- | --- | --- | --- |
|  |  | in WT | in BS | in WT | in BS | In WT | in BS |
| Alternative 3' of intron | Shorter exon | 47 | 29 | 8 | 49 | 8 | 15 |
| Alternative 5' of intron | Shorter exon | 34 | 29 | 7 | 46 | 14 | 14 |
| Mutually exclusive exon | Use of downstream exon | 27 | 31 | 65 | 38 | 5 | 10 |
| Retaining of an intron | Intron spliced out | 217 | 45 | 14 | 268 | 18 | 73 |
| Skipped exon | Exon excluded | 136 | 187 | 21 | 156 | 84 | 143 |
| Total | Any | 782 |  | 672 |  | 384 |  |

**Supplementary Table S2B.**
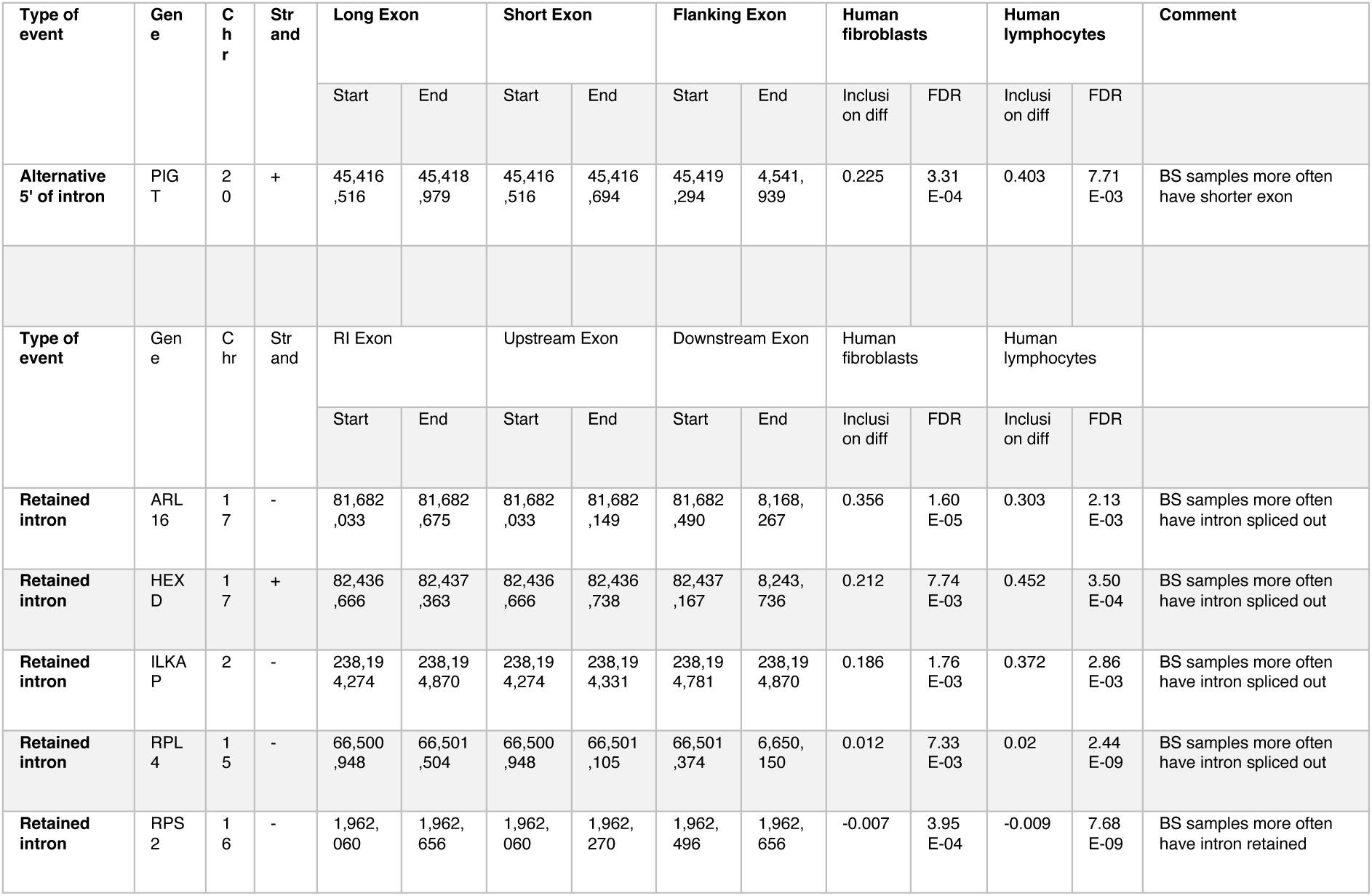
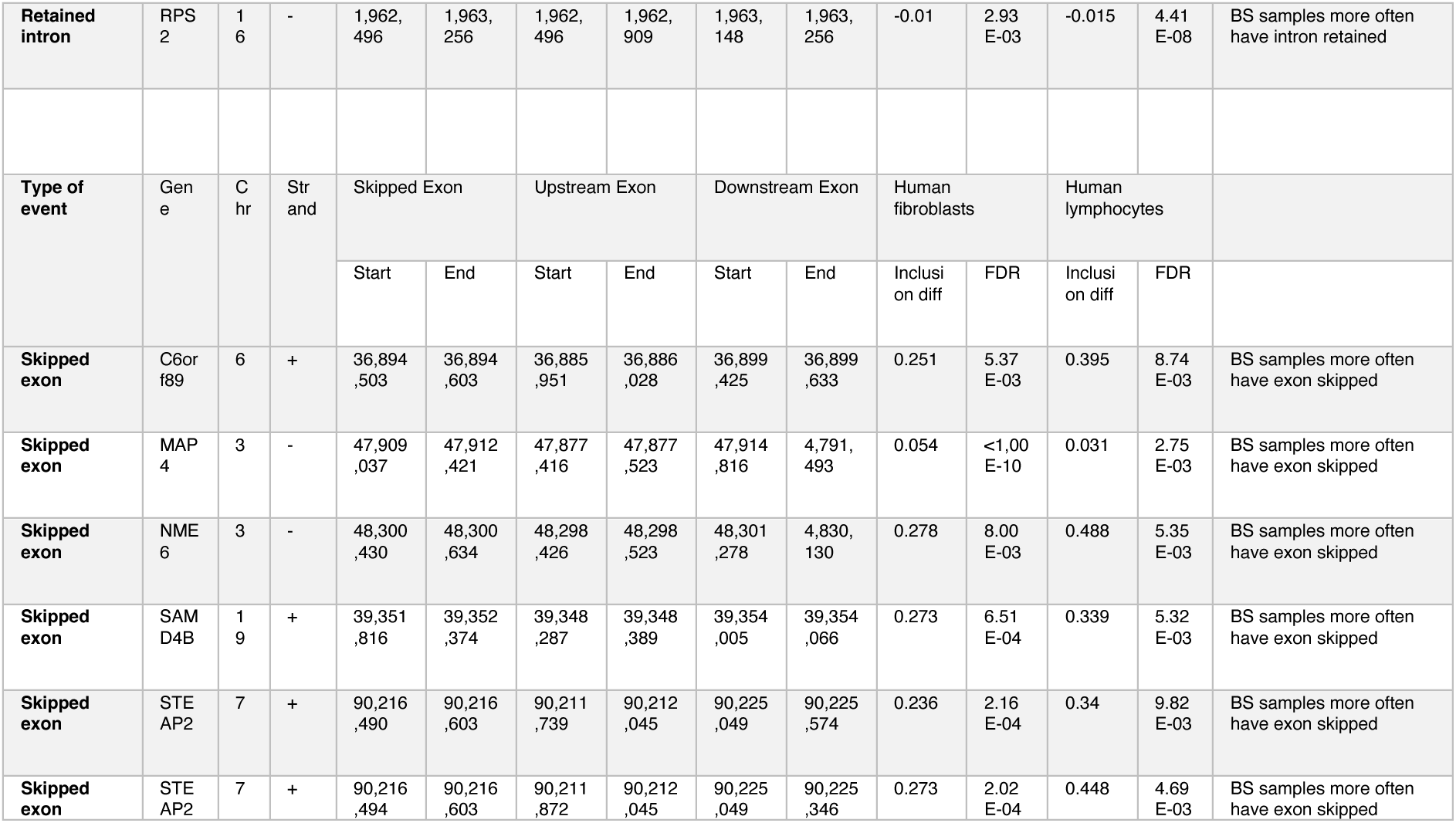
Alternative splicing events significant and having same direction between human fibroblast and lymphocyte datasets.

| Type of event | Gene | Chr | Strand | Long Exon |  | Short Exon |  | Flanking Exon |  | Human fibroblasts |  | Human lymphocytes |  | Comment |
| --- | --- | --- | --- | --- | --- | --- | --- | --- | --- | --- | --- | --- | --- | --- |
|  |  |  |  | Start | End | Start | End | Start | End | Inclusion diff | FDR | Inclusion diff | FDR |  |
| Alternative 5' of intron | PIGT | 20 | + | 45,416,516 | 45,418,979 | 45,416,516 | 45,416,694 | 45,419,294 | 4,541,939 | 0.225 | 3.31E-04 | 0.403 | 7.71E-03 | BS samples more often have shorter exon |
| Type of event | Gene | Chr | Strand | RI Exon |  | Upstream Exon |  | Downstream Exon |  | Human fibroblasts |  | Human lymphocytes |  | Comment |
|  |  |  |  | Start | End | Start | End | Start | End | Inclusion diff | FDR | Inclusion diff | FDR |  |
| Retained intron | ARL16 | 17 | - | 81,682,033 | 81,682,675 | 81,682,033 | 81,682,149 | 81,682,490 | 8,168,267 | 0.356 | 1.60E-05 | 0.303 | 2.13E-03 | BS samples more often have intron spliced out |
| Retained intron | HEXD | 17 | + | 82,436,666 | 82,437,363 | 82,436,666 | 82,436,738 | 82,437,167 | 8,243,736 | 0.212 | 7.74E-03 | 0.452 | 3.50E-04 | BS samples more often have intron spliced out |
| Retained intron | ILKAP | 2 | - | 238,194,274 | 238,194,870 | 238,194,274 | 238,194,331 | 238,194,781 | 238,194,870 | 0.186 | 1.76E-03 | 0.372 | 2.86E-03 | BS samples more often have intron spliced out |
| Retained intron | RPL4 | 15 | - | 66,500,948 | 66,501,504 | 66,500,948 | 66,501,105 | 66,501,374 | 6,650,150 | 0.012 | 7.33E-03 | 0.02 | 2.44E-09 | BS samples more often have intron spliced out |
| Retained intron | RPS2 | 16 | - | 1,962,060 | 1,962,656 | 1,962,060 | 1,962,270 | 1,962,496 | 1,962,656 | -0.007 | 3.95E-04 | -0.009 | 7.68E-09 | BS samples more often have intron retained |
| Retained intron | RPS 2 | 16 | - | 1,962,496 | 1,963,256 | 1,962,496 | 1,962,909 | 1,963,148 | 1,963,256 | -0.01 | 2.93 E-03 | -0.015 | 4.41 E-08 | BS samples more often have intron retained |
| Type of event | Gene | Chr | Strand | Skipped Exon |  | Upstream Exon |  | Downstream Exon |  | Human fibroblasts |  | Human lymphocytes |  |  |
|  |  |  |  | Start | End | Start | End | Start | End | Inclusion diff | FDR | Inclusion diff | FDR |  |
| Skipped exon | C6orf89 | 6 | + | 36,894,503 | 36,894,603 | 36,885,951 | 36,886,028 | 36,899,425 | 36,899,633 | 0.251 | 5.37 E-03 | 0.395 | 8.74 E-03 | BS samples more often have exon skipped |
| Skipped exon | MAP4 | 3 | - | 47,909,037 | 47,912,421 | 47,877,416 | 47,877,523 | 47,914,816 | 4,791,493 | 0.054 | <1,00 E-10 | 0.031 | 2.75 E-03 | BS samples more often have exon skipped |
| Skipped exon | NME6 | 3 | - | 48,300,430 | 48,300,634 | 48,298,426 | 48,298,523 | 48,301,278 | 4,830,130 | 0.278 | 8.00 E-03 | 0.488 | 5.35 E-03 | BS samples more often have exon skipped |
| Skipped exon | SAMD4B | 19 | + | 39,351,816 | 39,352,374 | 39,348,287 | 39,348,389 | 39,354,005 | 39,354,066 | 0.273 | 6.51 E-04 | 0.339 | 5.32 E-03 | BS samples more often have exon skipped |
| Skipped exon | STEAP2 | 7 | + | 90,216,490 | 90,216,603 | 90,211,739 | 90,212,045 | 90,225,049 | 90,225,574 | 0.236 | 2.16 E-04 | 0.34 | 9.82 E-03 | BS samples more often have exon skipped |
| Skipped exon | STEAP2 | 7 | + | 90,216,494 | 90,216,603 | 90,211,872 | 90,212,045 | 90,225,049 | 90,225,346 | 0.273 | 2.02 E-04 | 0.448 | 4.69 E-03 | BS samples more often have exon skipped |

**Supplementary Table S3.** Analysis of G4 overrepresentation in intronic flanks (250bp 5’of intron) of skipped exons.

| Human fibroblasts |  | Positive, **skipped** in BS |  |  |  | Negative, **retained** In BS |  |  |  |
| --- | --- | --- | --- | --- | --- | --- | --- | --- | --- |
|  |  | Significant AS | Insignificant | Significant. % | P-value | Significant AS | Insignificant | Significant. % | P-value |
| Upstream exon | G4 on coding strand | 28 | 2695 | 1.03 | 0.661 | 15 | 2463 | 0.61 | 0.645 |
|  | G4 on template strand | 22 | 1309 | 1.65 | 6.43E-03 | 10 | 1205 | 0.82 | 0.520 |
| Skipped exon | G4 on coding strand | 3 | 1071 | 0.28 | 1.95E-02 | 8 | 955 | 0.83 | 0.549 |
|  | G4 on template strand | 8 | 713 | 1.11 | 0.658 | 7 | 662 | 1.05 | 0.235 |
| Downstream exon | G4 on coding strand | 8 | 790 | 1.00 | 0.882 | 4 | 845 | 0.47 | 0.456 |
|  | G4 on template strand | 7 | 504 | 1.37 | 0.325 | 4 | 540 | 0.74 | 0.865 |
| Total irrespective of G4 presence |  | 187 | 19447 | 0.95 |  | 136 | 19971 | 0.68 | 2.18E-03 |
| Human lymphocytes |  | Positive, **skipped** in BS |  |  |  | Negative, **retained** In BS |  |  |  |
|  |  | Significant AS | Insignificant | Significant. % | P-value | Significant AS | Insignificant | Significant. % | P-value |
| Upstream exon | G4 on coding strand | 29 | 3120 | 0.92 | 3.15E-02 | 2 | 1703 | 0.12 | 0.793 |
|  | G4 on template strand | 9 | 1462 | 0.61 | 0.903 | 2 | 891 | 0.22 | 0.486 |
| Skipped exon | G4 on coding strand | 6 | 1247 | 0.48 | 0.472 | 1 | 797 | 0.13 | 0.912 |
|  | G4 on template strand | 5 | 854 | 0.58 | 0.839 | 1 | 546 | 0.18 | 0.783 |
| Downstream exon | G4 on coding strand | 5 | 964 | 0.52 | 0.631 | 1 | 635 | 0.16 | 0.903 |
|  | G4 on template strand | 2 | 607 | 0.33 | 0.333 | 0 | 407 | 0.00 | 0.444 |
| Total irrespective of G4 presence |  | 156 | 24359 | 0.64 |  | 21 | 15021 | 0.14 | 6.68E-13 |

| Mouse ES cells |  | Positive, **skipped** in BS |  |  |  | Negative, **retained** In BS |  |  |  |
| --- | --- | --- | --- | --- | --- | --- | --- | --- | --- |
|  |  | Significant AS | Insignificant | Significant. % | P-value | Significant AS | Insignificant | Significant. % | P-value |
| **Upstream exon** | G4 on coding strand | 9 | 1186 | 0.75 | 0.377 | 6 | 910 | 0.66 | 0.753 |
|  | G4 on template strand | 8 | 606 | 1.30 | 0.434 | 3 | 517 | 0.58 | 0.656 |
| **Skipped exon** | G4 on coding strand | 6 | 589 | 1.01 | 0.976 | 2 | 424 | 0.47 | 0.506 |
|  | G4 on template strand | 1 | 407 | 0.25 | 0.121 | 2 | 335 | 0.59 | 0.749 |
| **Downstream exon** | G4 on coding strand | 3 | 528 | 0.56 | 0.308 | 0 | 389 | 0.00 | 0.083 |
|  | G4 on template strand | 2 | 364 | 0.55 | 0.380 | 5 | 317 | 1.55 | 0.085 |
| **Total irrespective of G4 presence** |  | 143 | 14210 | 1.00 |  | 84 | 11258 | 0.74 | 2.96E-02 |

**Supplementary Table S4.** Clustering of differentially regulated genes in genome.

| Gene 1 direction | Gene 2 direction | Strands | Human fibroblasts | Human lymphocytes | Mouse ES cells |
| --- | --- | --- | --- | --- | --- |
| Up | Up | Same | 265 | 10 | 4 |
| Up | Up | Opposite | 294 | 16 | 8 |
| Up | Down | Same | 118 | 1 | 0 |
| Up | Down | Opposite | 141 | 3 | 3 |
| Down | Up | Same | 125 | 2 | 3 |
| Down | Up | Opposite | 144 | 2 | 1 |
| Down | Down | Same | 229 | 56 | 11 |
| Down | Down | Opposite | 248 | 45 | 12 |
| **Total expected same direction** |  |  | 784 | 84 | 21 |
| **Total expected opposite direction** |  |  | 780 | 51 | 21 |
| **Total observed same direction** |  |  | 1036 | 127 | 35 |
| **Total observed opposite direction** |  |  | 528 | 8 | 7 |

**Supplementary Table S5A.**
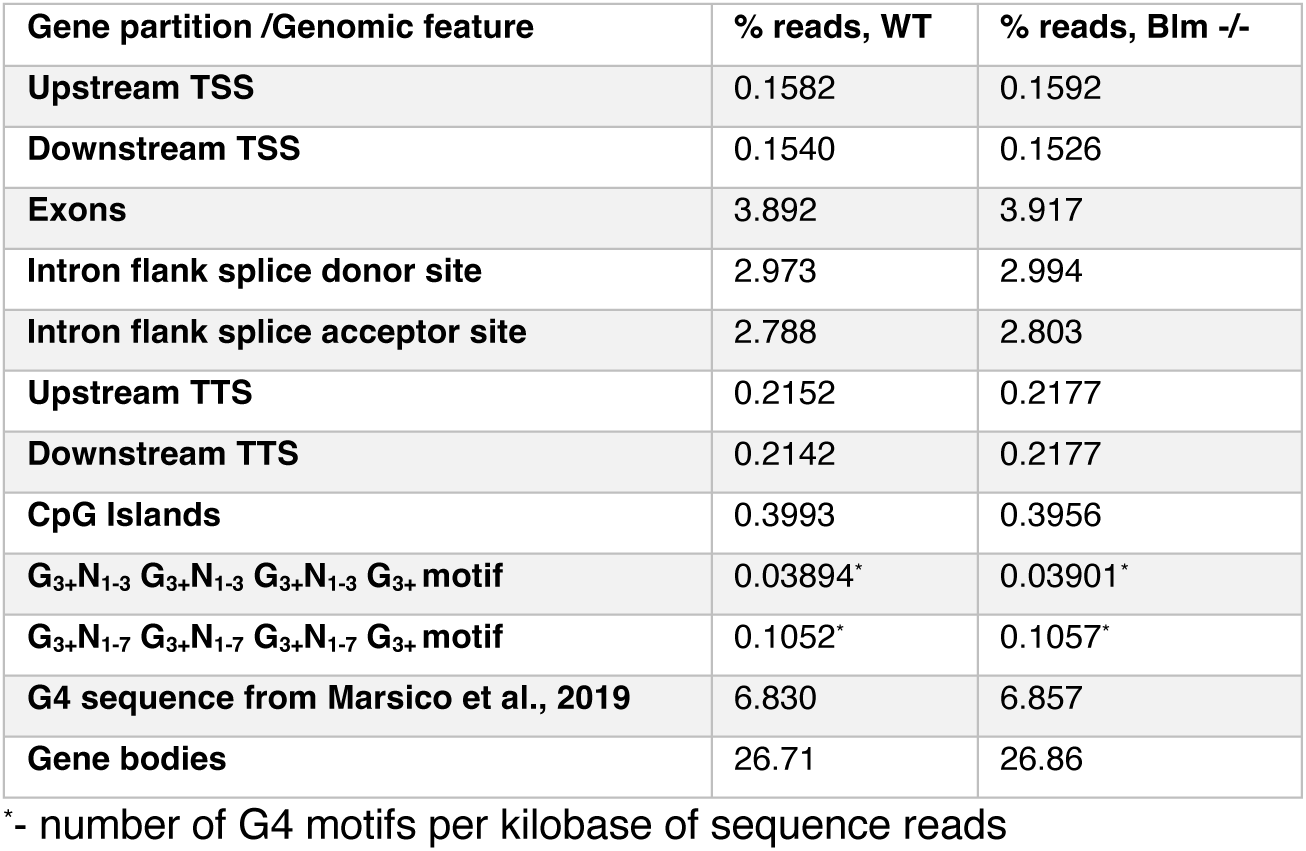
Nucleosome occupancy at different gene partitions in WT and Blm-/– mouse ES cells.

| Gene partition /Genomic feature | % reads, WT | % reads, Blm<sup>-/-</sup> |
| --- | --- | --- |
| **Upstream TSS** | 0.1582 | 0.1592 |
| **Downstream TSS** | 0.1540 | 0.1526 |
| **Exons** | 3.892 | 3.917 |
| **Intron flank splice donor site** | 2.973 | 2.994 |
| **Intron flank splice acceptor site** | 2.788 | 2.803 |
| **Upstream TTS** | 0.2152 | 0.2177 |
| **Downstream TTS** | 0.2142 | 0.2177 |
| **CpG Islands** | 0.3993 | 0.3956 |
| **G<sub>3+</sub>N<sub>1-3</sub> G<sub>3+</sub>N<sub>1-3</sub> G<sub>3+</sub>N<sub>1-3</sub> G<sub>3+</sub> motif** | 0.03894* | 0.03901* |
| **G<sub>3+</sub>N<sub>1-7</sub> G<sub>3+</sub>N<sub>1-7</sub> G<sub>3+</sub>N<sub>1-7</sub> G<sub>3+</sub> motif** | 0.1052* | 0.1057* |
| **G4 sequence from Marsico et al., 2019** | 6.830 | 6.857 |
| **Gene bodies** | 26.71 | 26.86 |

\*- number of G4 motifs per kilobase of sequence reads

**Supplementary Table S5B.**
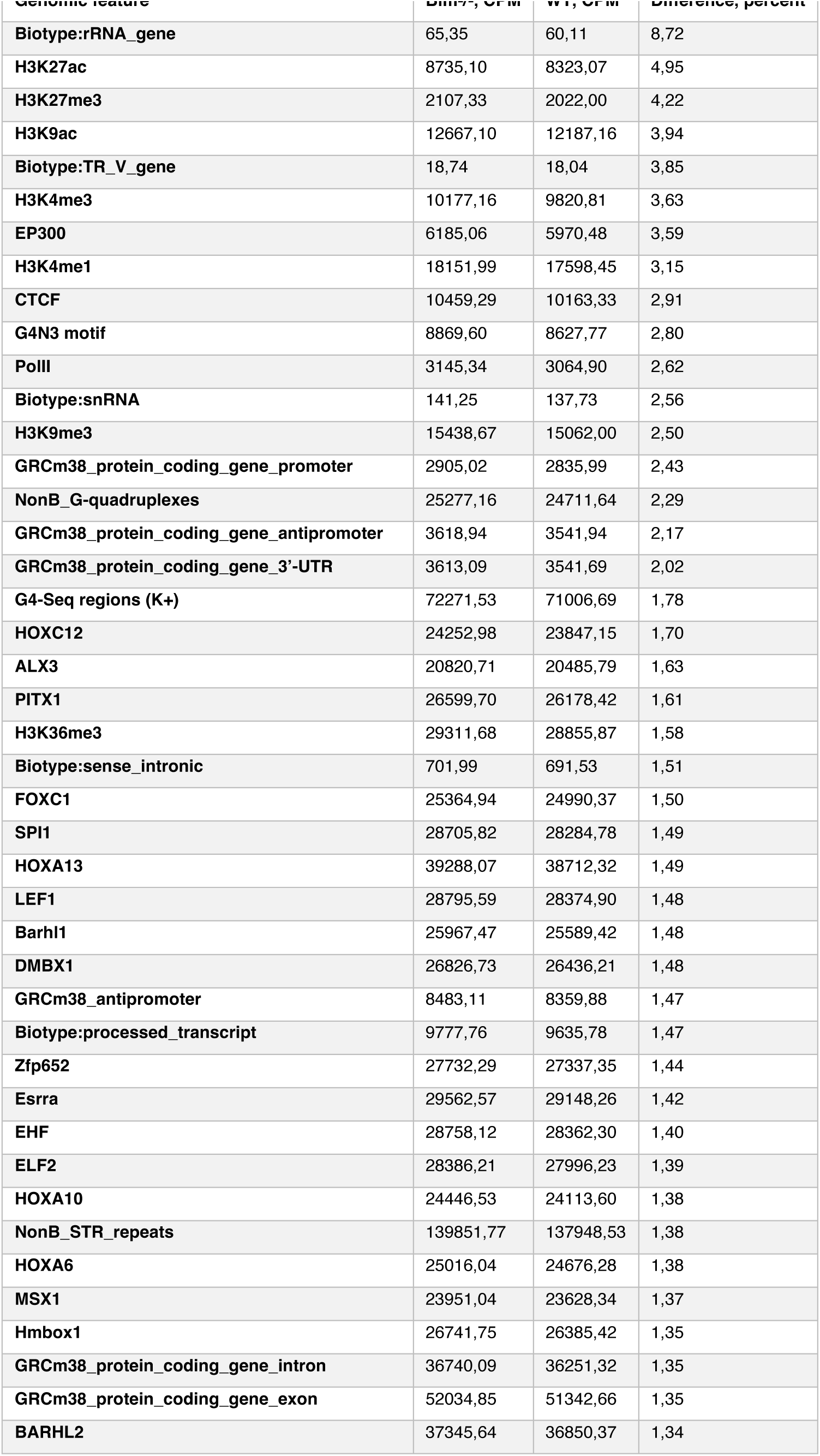
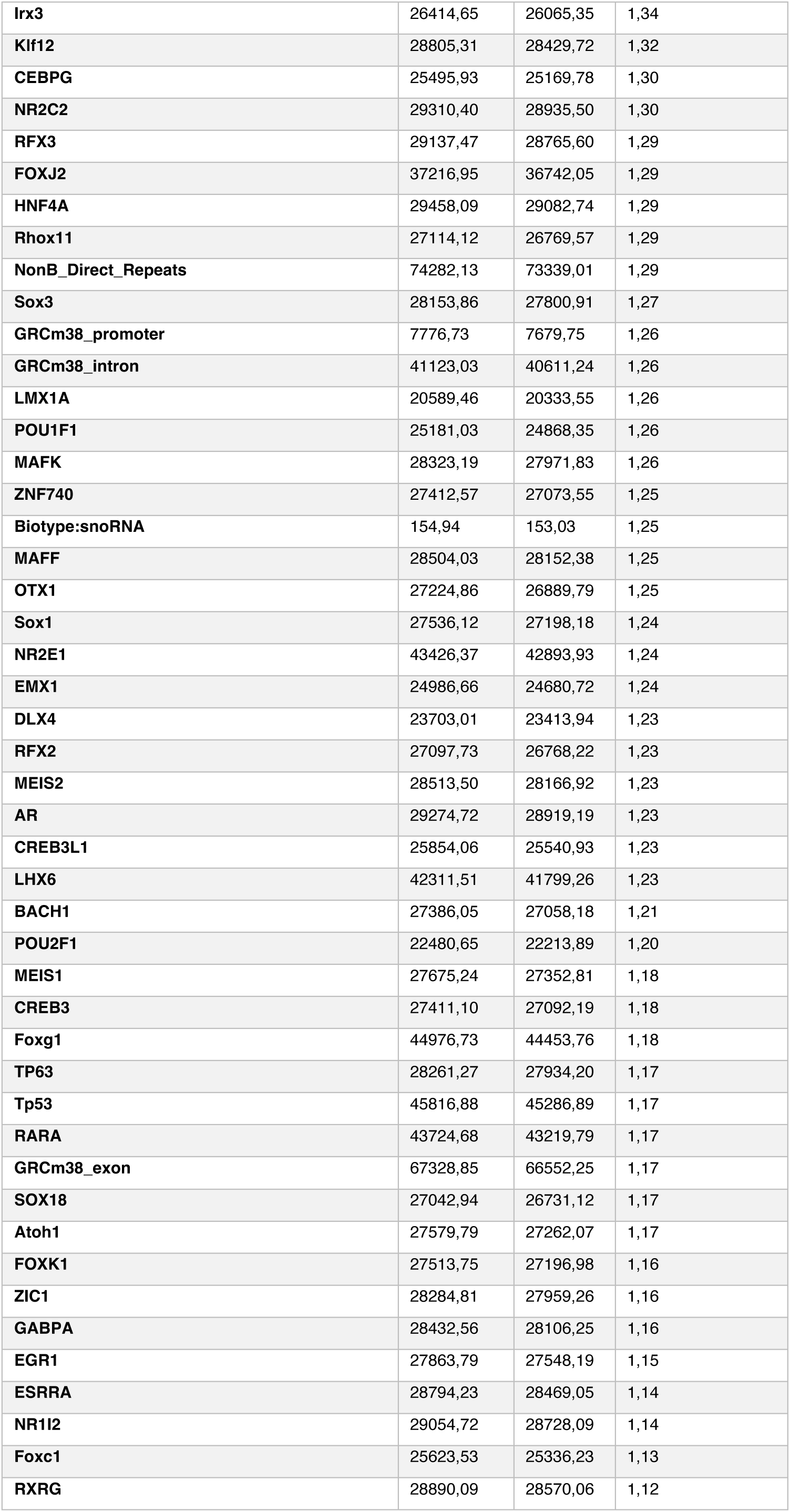
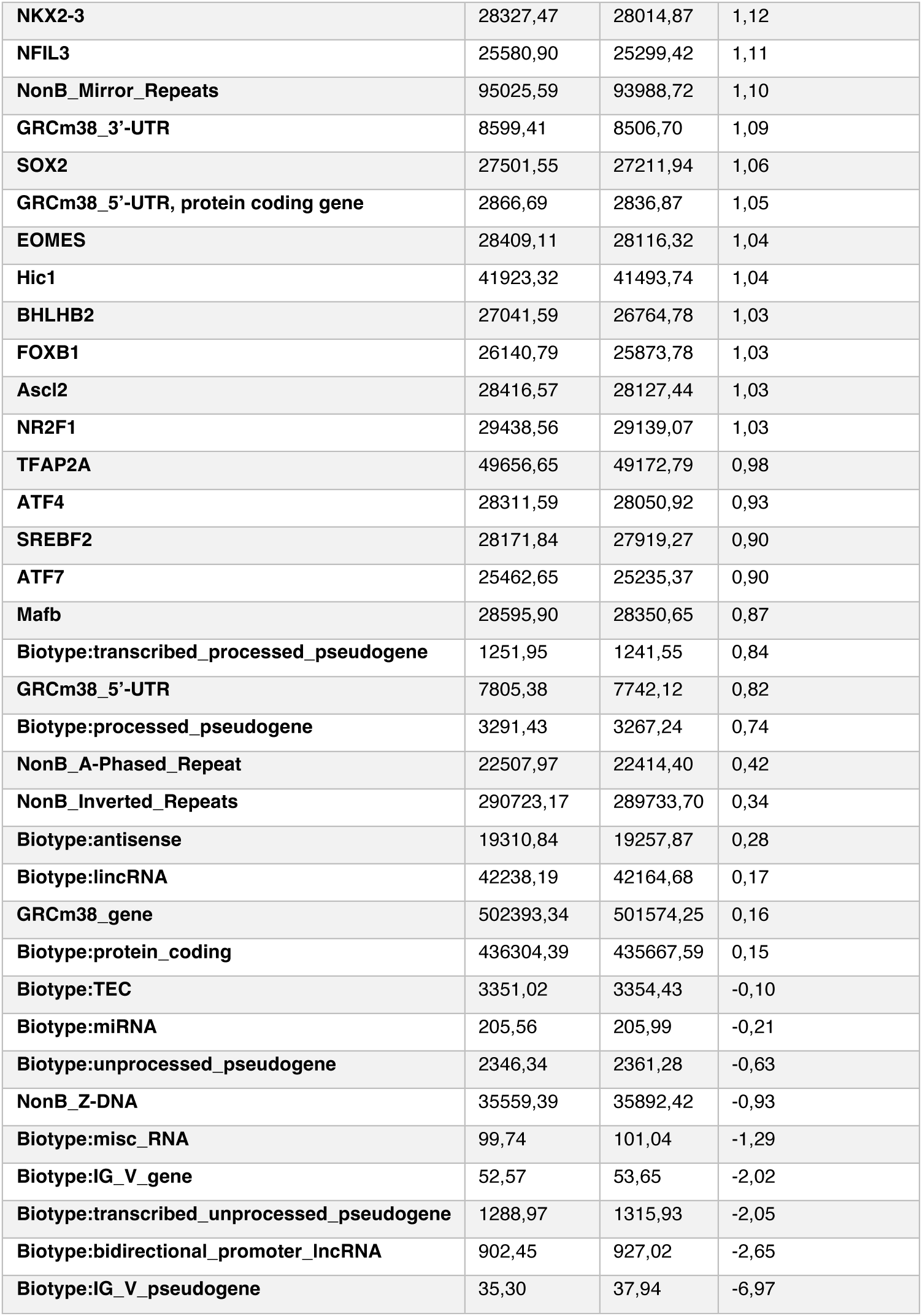
Nucleosome occupancy at different genomic features in WT and Blm-/– mouse ES cells.

**Supplementary Table S6.** Top 10 enriched GO terms in differentially expressed gene sets (FDR = 0.10) in human fibroblasts (microarray data)

| Microarray data: human fibroblasts |  |  |  |
| --- | --- | --- | --- |
| Top 10 enriched GO terms in upregulated genes (n=199, FDR= 0.10) |  |  |  |
| Id | Term | Ont | P-value |
| GO:0060255 | regulation of macromolecule metabolic process | BP | 2.7456E-07 |
| GO:0098772 | molecular function regulator | MF | 5.063E-07 |
| **GO:0080090** | regulation of primary metabolic process | BP | 5.4517E-07 |
| **GO:0051171** | regulation of nitrogen compound metabolic process | BP | 9.0247E-07 |
| **GO:0019222** | regulation of metabolic process | BP | 1.5015E-06 |
| **GO:0031323** | regulation of cellular metabolic process | BP | 1.5823E-06 |
| **GO:0048522** | positive regulation of cellular process | BP | 4.8267E-06 |
| **GO:0050794** | regulation of cellular process | BP | 5.8057E-06 |
| **GO:0005173** | stem cell factor receptor binding | MF | 7.8833E-06 |
| **GO:1902667** | regulation of axon guidance | BP | 1.0147E-05 |
| **Top 10 enriched GO terms in downregulated genes (n=56, FDR= 0.10)** |  |  |  |
| **Id** | **Term** | **Ont** | **P-value** |
| **GO:0006936** | muscle contraction | BP | 1.5859E-05 |
| **GO:0061061** | muscle structure development | BP | 3.1451E-05 |
| **GO:0003012** | muscle system process | BP | 7.7716E-05 |
| **GO:0005198** | structural molecule activity | MF | 0.00013975 |
| **GO:0030240** | skeletal muscle thin filament assembly | BP | 0.0001847 |
| **GO:0007517** | muscle organ development | BP | 0.00019947 |
| **GO:0033275** | actin-myosin filament sliding | BP | 0.00030398 |
| **GO:0030049** | muscle filament sliding | BP | 0.00030398 |
| **GO:0014866** | skeletal myofibril assembly | BP | 0.0003432 |
| **GO:0003008** | system process | BP | 0.00058819 |

**Supplementary Table S7.**
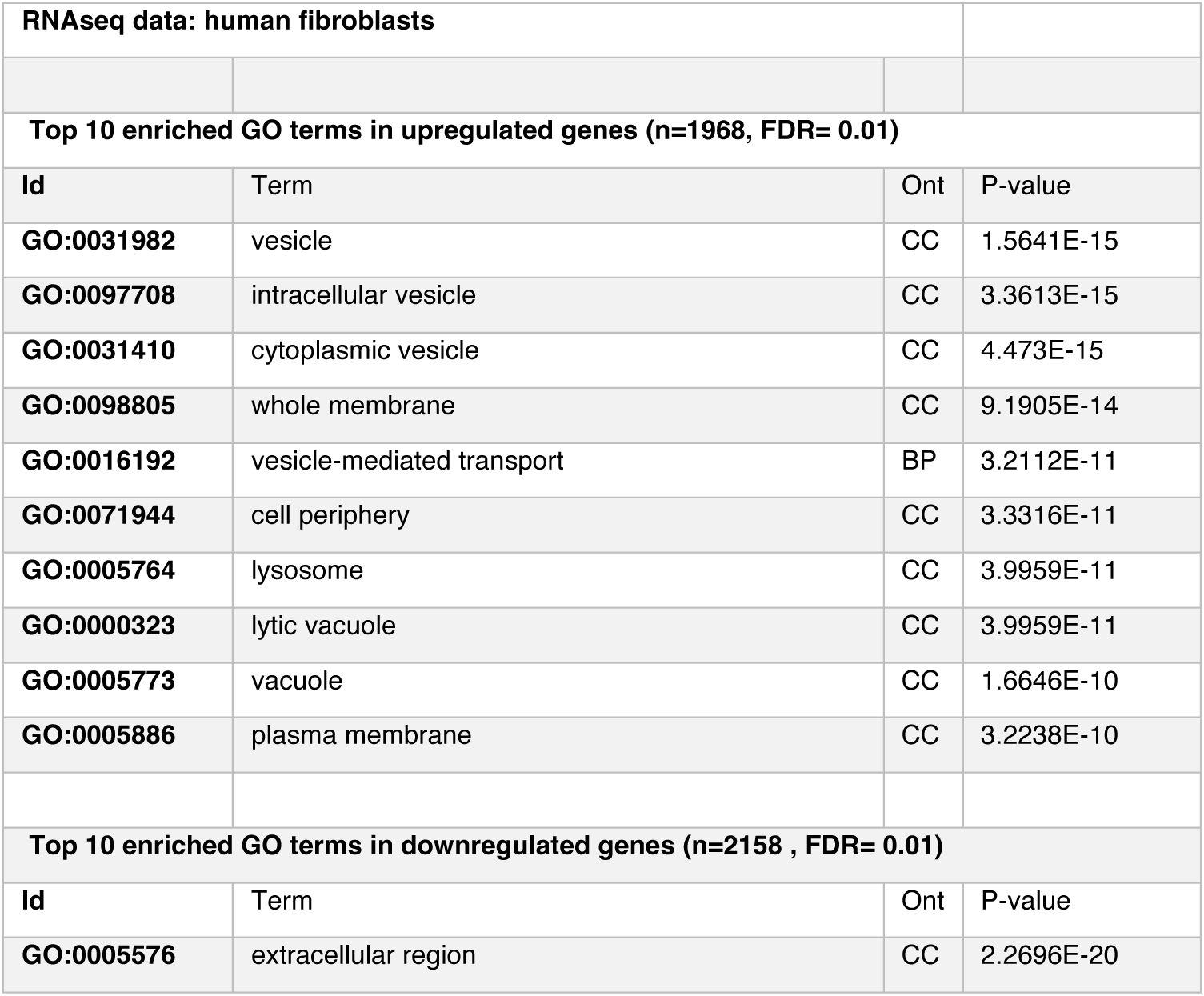
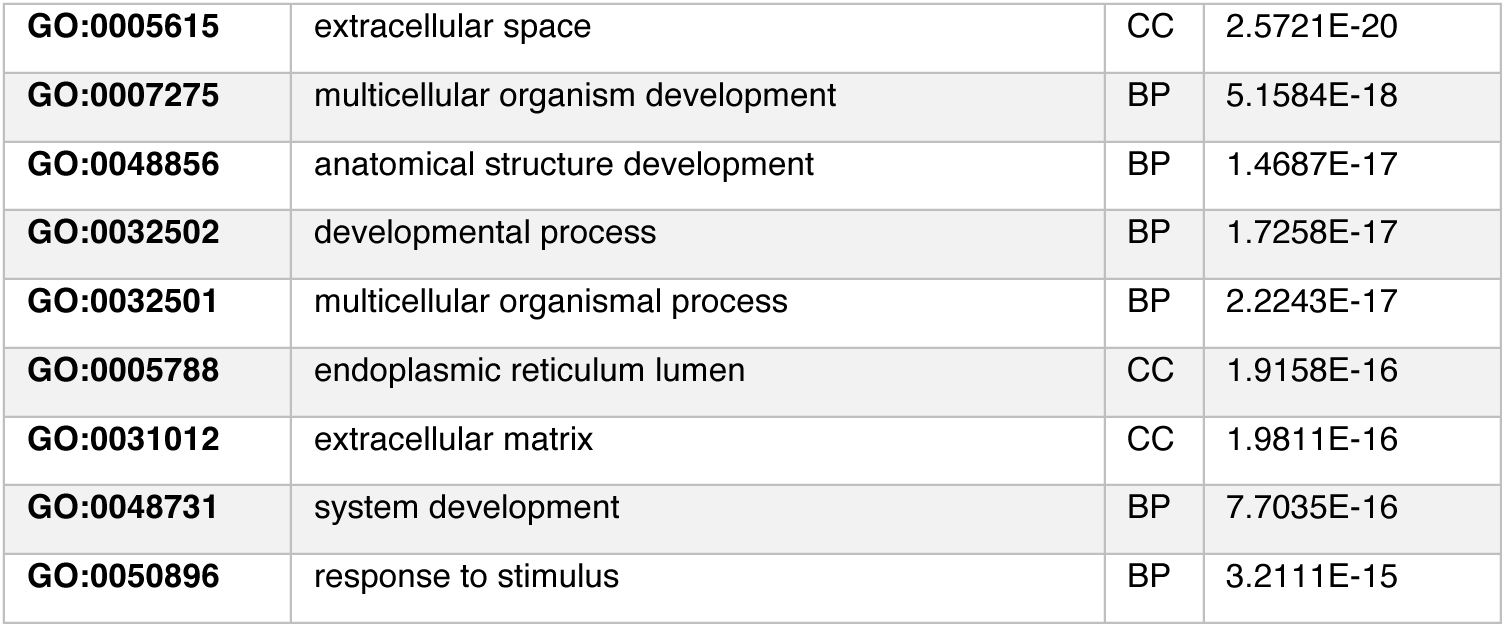
Top 10 enriched GO terms in differentially expressed gene sets (FDR = 0.01) in human fibroblasts (RNAseq data)

**Supplementary Table S8.** Top 10 enriched GO terms in differentially expressed gene sets (FDR = 0.01) in human lymphocytes (RNAseq data)

| RNAseq data: human lymphocytes |  |  |  |
| --- | --- | --- | --- |
| **Top 10 enriched GO terms in upregulated genes (n=204, FDR= 0.01)** |  |  |  |
| Id | Term | Ont | P-value |
| **GO:0098798** | mitochondrial protein complex | CC | 2.9804E-09 |
| **GO:0098800** | inner mitochondrial membrane protein complex | CC | 6.0201E-09 |
| **GO:0019866** | organelle inner membrane | CC | 1.9241E-08 |
| **GO:0005743** | mitochondrial inner membrane | CC | 2.171E-08 |
| **GO:0006396** | RNA processing | BP | 6.0796E-08 |
| **GO:0098803** | respiratory chain complex | CC | 8.9898E-08 |
| **GO:0005746** | mitochondrial respirasome | CC | 1.8555E-07 |
| **GO:0031966** | mitochondrial membrane | CC | 3.2985E-07 |
| **GO:0070469** | respirasome | CC | 3.9704E-07 |
| **GO:0005740** | mitochondrial envelope | CC | 9.2092E-07 |
| **Top 10 enriched GO terms in downregulated genes (n=598 , FDR= 0.01)** |  |  |  |
| Id | Term | Ont | P-value |
| **GO:0062023** | collagen-containing extracellular matrix | CC | 0.00016379 |
| **GO:0005132** | type I interferon receptor binding | MF | 0.00021847 |
| **GO:0031012** | extracellular matrix | CC | 0.00048964 |
| **GO:0097676** | histone H3-K36 dimethylation | BP | 0.000649 |
| **GO:0033141** | positive regulation of peptidyl-serine phosphorylation of STAT protein | BP | 0.000649 |
| **GO:0097503** | sialylation | BP | 0.00082099 |
| **GO:0008373** | sialyltransferase activity | MF | 0.00082099 |
| **GO:0030021** | extracellular matrix structural constituent conferring compression resistance | MF | 0.00128532 |
| **GO:0006895** | Golgi to endosome transport | BP | 0.00186865 |
| **GO:0045053** | protein retention in Golgi apparatus | BP | 0.00212131 |

**Supplementary Table S9.**
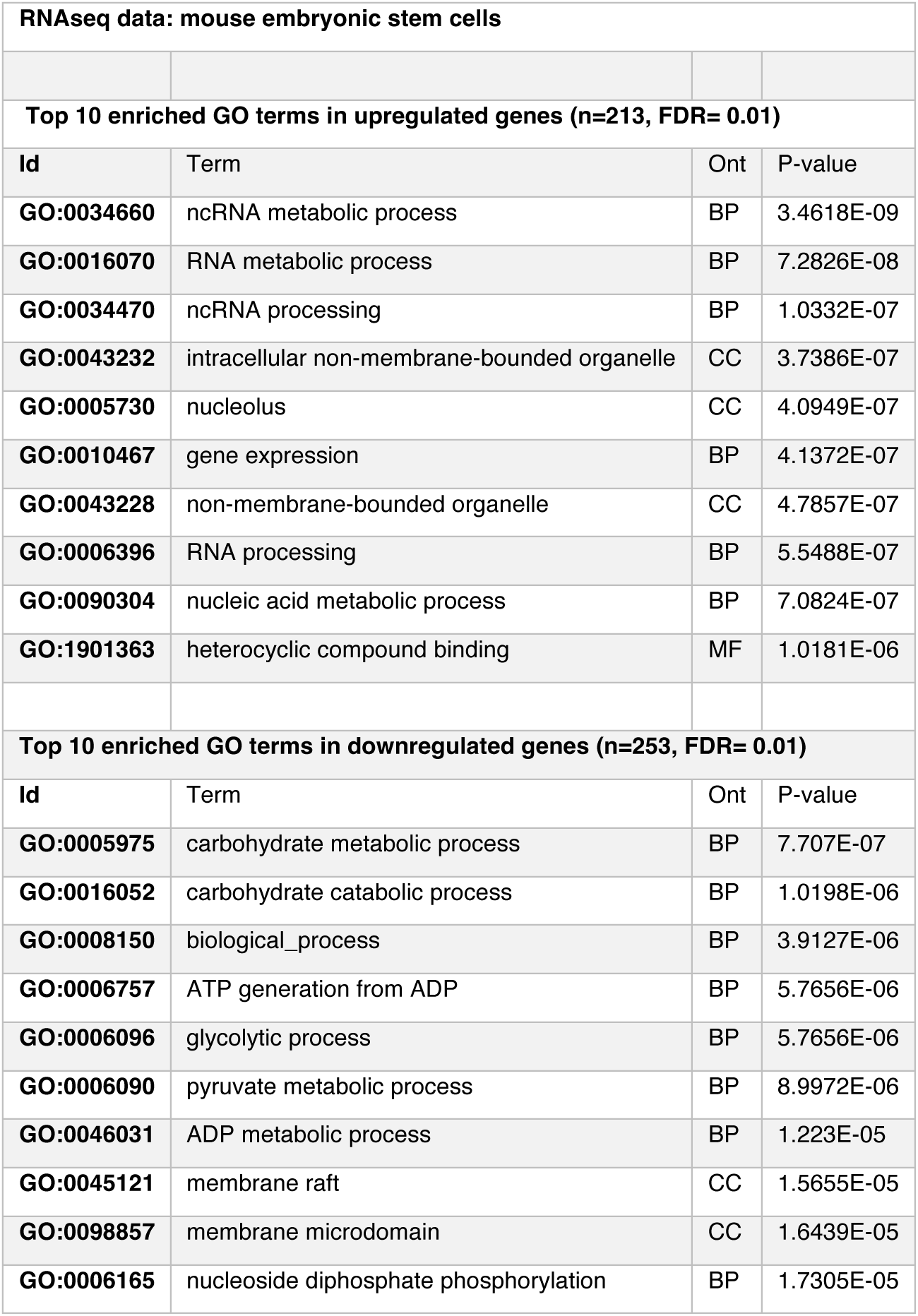
Top 10 enriched GO terms in differentially expressed gene sets (FDR = 0.01) in mouse embryonic stem cells (RNAseq data)

